# Mitochondrial Dysfunction Induces Epigenetic Dysregulation by H3K27 Hyperacetylation to Perturb Active Enhancers in Parkinson’s Disease Models

**DOI:** 10.1101/808246

**Authors:** Minhong Huang, Dan Lou, Adhithiya Charli, Dehui Kong, Huajun Jin, Vellareddy Anantharam, Arthi Kanthasamy, Zhibin Wang, Anumantha G. Kanthasamy

**Author notes:** Both authors contributed equally to this work (Minhong Huang, Dan Lou). To whom correspondence should be addressed: Anumantha Kanthasamy, Ph.D., Distinguished Professor and Chair, Department of Biomedical Sciences, Iowa State University, Ames, IA 50011., Zhibin Wang, Ph.D., Associate Professor, Laboratory of Environmental Epigenomes, Department of Environmental Health and Engineering, Bloomberg School of Public Health, Johns Hopkins University, Baltimore, MD 21205.

## Abstract

Genetic mutations explain only 10-15% of cases of Parkinson’s disease (PD), while an overriding environmental component has been implicated in the etiopathogenesis of PD. But regardless of where the underlying triggers for the onset of familial and sporadic PD fall on the gene-environment axis, mitochondrial dysfunction emerges as a common mediator of dopaminergic neuronal degeneration. Herein, we employ a multidisciplinary approach to convincingly demonstrate that neurotoxicant exposure- and genetic mutation-driven mitochondrial dysfunction share a common mechanism of epigenetic dysregulation. Under both scenarios, lysine 27 acetylation of likely variant H3.2 (H3.2K27ac) increased in dopaminergic neuronal models of PD, thereby opening that region to active enhancer activity via H3K27 hyperacetylation. These vulnerable epigenomic loci represent potential transcription factor motifs for PD pathogenesis. We further confirmed the mitochondrial dysfunction induced H3K27ac during neurodegeneration in *ex vivo* models of PD. Our results reveal an exciting axis of ‘exposure/mutation-mitochondrial dysfunction-metabolism-H3K27ac-transcriptome’ for PD pathogenesis. Collectively, the novel mechanistic insights presented here interlinks mitochondrial dysfunction to epigenetic transcriptional regulation in dopaminergic degeneration as well as offer potential new epigenetic intervention strategies for PD.

## Introduction

Since the first reports of *SNCA* mutation in the pathogenesis of Parkinson’s disease (PD) (Polymeropoulos et al., 1997), investigations on genetic causes in PD have flourished and gained mainstream attention in the scientific community. However, genetic mutations at 15 loci only explain about 10-15% of PD cases (Dawson et al., 2010; Goldman, 2014). Notably, mutations of several key PD genes, including *PARK2* (Parkin), *PARK6* (PINK1), *SNCA*, and *LRRK2*, modify key mitochondrial functions (Trancikova et al., 2012; Zanon et al., 2018). On the other hand, environmental exposure and its interactions with genetic factors are expected to explain the majority of PD cases. Many environmental neurotoxicants are known to impair mitochondrial functions. For example, the mitochondrial complex-I inhibitor rotenone has been implicated in the etiology of PD (Betarbet et al., 2006; Betarbet et al., 2000; Charli et al., 2016; Hatcher et al., 2008; Tanner et al., 2011; Wang et al., 2006). The neurotoxicity of rotenone exposure is correlated with increased oxidative stress, a damaged ubiquitin-proteasome system (UPS), mitochondrial dysfunction, and impaired neurotransmitter systems (Tanner et al., 2011). Rotenone inhibits healthy mitochondrial functionality by impairing oxidative phosphorylation and protein synthesis, leading to energy depletion (Li et al., 2003; Wu et al., 2018). Collectively, mitochndrial dysfunction is key for both genetically and environmentally linked PD; however, the underlying mechanism is not clear.

For non-mutagenic pathways, such as neurotoxicant exposure, PD pathogenesis is expected through epigenetic mechanisms. Recent studies indeed imply that epigenomic alteration plays an important role in the neuropathology and etiology of PD (Horgusluoglu et al., 2017). We previously discovered that rotenone-induced mitochondrial disorders and accumulation of the major histone acetyltransferase enzyme CBP (CREB-binding protein) led to aberrant increases in the acetylation of core histones in dopamin(DA)ergic cell culture (Song et al., 2010). Increased acetylation of core histones in an animal model of PD and in human PD brains has been reported (Harrison et al., 2018). Hyperacetylation makes neurons vulnerable to environmental toxicity, thereby increasing the risk of developing PD (Song et al., 2010). Though epigenetic modifications have triggered researchers’ interest in the context of neurodegneration, the detailed mechanism of neurotoxic pesticide-induced histone modification and its core role in environmentally related PD remains unclear.

Among many histone acetylation (HAc) marks, H3K27ac is a robust mark of active promoters and distal enhancers that are tightly coupled to gene expression and transcription factor (TF)-binding, and as a cis-regulatory element, is conservative across different cell types (Heintzman et al., 2009; Marzi et al., 2018; Wang et al., 2008). Intriguingly, a recent publication also found that some H3K27ac-marked enhancers (that is, latent enhancers) acquired histone marks in response to stimulation and enabled H3K27ac deposition, even though they did not have such characterisitic before stimulus-evoked activation (Ostuni et al., 2013). Therefore, by marking the genome, H3K27ac acts as an epigenomic memory of environmental stimuli and mediates a stronger response upon subsequent stimulation. It represents a homeostatic mechanism that guarantees flexibility and adaptability to environmental change (Haley et al., 2019). Aberrant H3K27ac modification triggered by environmental stimuli would consequently induce pervasive changes in gene expression. Alterations in gene expression are considered as a fundamental hallmark of environmentally related PD (Borrageiro et al., 2018; Li et al., 2019; Valentine et al., 2019). Comprehensive genome-wide analysis of gene expression alterations and H3K27ac modifications can better elucidate the complex cooperative biological processes occurring in the genome during PD pathogenesis. However, such high-resolution analysis has never been performed in PD models, and the pattern by which H3K27ac modification cooperates with environmental stimuli and mitochondria impairment to affect PD pathogenesis remains to be elucidated.

Herein, we aim to further explore the predominant mitochondrial dysfunctions in PD, with a focus on how altering the histone code contributes to PD pathogenesis. To uncover molecular mechanisms in nigral DAergic neuronal degeneration following exposure to a mitochondria-impairing neurotoxicant, we mapped the acetylation profile across the genome with a focus on core histone H3. We characterized H3K27 acetylation modification and mRNA transcription levels at certain genomic loci in multiple PD models, including DAergic neuronal cell culture model of PD and *ex vivo* nigral brain slice cultureWe performed genome-wide ChIP-Seq characterization of H3K27ac as well as transcriptome profiling in DAergic neurons whose mitochondria had been impaired either by exposure to a neurotoxic pesticide or by genetic modification. Integrated bioinformatics analysis of ChIP-Seq and RNA-Seq data were performed to unravel the potential functional role of H3K27ac modification in PD progression. To the best of our knowledge, this is the first time the vulnerable epigenomic loci regulated by H3K27 hyperacetylation in environmental neurotoxicant-induced dopaminergic neurodegeneration have been identified.

## Results

### Genome-wide hyperacetylation profiling identifies H3K27ac as the key lysine acetylation site in mitochondria-impaired nigral DAergic neurons

Mitochondrial dysfunction has been linked to the alteration of the histone post-translocation in several disease conditions, but it has not been explored in detail in PD. With mitochondrial dysfunction predominant in many familial PD cases and almost all environmental linked PD. We 1) explored the possibility that even more severe changes in HAc codes are involved in PD pathogenesis, and 2) searched for epigenomic loci that are vulnerable to neurotoxicant-induced alterations in HAc, as implicated by our pioneering report (Song et al., 2010). In our efforts to untangle the underlying mechanism of hyperacetylation and to map acetylation sites that contribute to neurodegeneration, this is, to our knowledge, the first evidence linking HAc to environmental pesticide-induced PD.

To deal with the low abundance and substoichiometric nature of potential changes (Baeza et al., 2014; Choudhary et al., 2014; Weinert et al., 2014), we used an optimized AcetylScan method that combines antibody enrichment of PTM-containing peptides with LC-MS/MS to map the lysine acetylation (Kac) sites of the core histones H3/H4. We exposed nigral DAergic N27 neuronal cells to the mitochondrial complex I inhibitor rotenone in a low-dose exposure paradigm (1 μM rotenone for 3 h). Cells were processed for Kac sites detection according to Song et al. (2010) and Svinkina et al. (2015). Although the rotenone exposure was sufficient to inhibit mitochondrial complex-1, it did not induce significant cell death, as measured by MTS assay (Fig. S1). To confirm successful treatment, we probed the DNA damage biomarker γ-H2AX, as reported by Sánchez-Flores et al. (2015). Notably increased γ-H2AX occurred in rotenone-exposed samples as revealed by AcetylScan analysis, validating rotenone-induced DNA damage (Fig. 1A-B). From our novel AcetylScan method, we attained global, deep coverage of the acetylome, including 3171 acetylation peptides. The relative abundance of Kac peptide data was transformed to log2 for better statistical comparison and followed a median offset correction (normalization), which was set based on overall relative abundance values. Among 3165 (out of 3171) Kac peptides whose fold changes were larger than 2.5-fold, those of histones H3/H4 exhibited remarkably higher fold change in mitochondria-impaired N27 neurons than in naïve neurons. Consistent with our previous data (Song et al., 2010) as well as published papers, these results suggest PD-related mitochondrial impairment induces H3/H4 hyperacetylation. Of at least 10 possible Kac sites on H3 (K4, K9, K14, K18, K23, K27, K36, K56, K64, and K79) and four on H4 (K5, K8, K12, and K16) (Garcia et al., 2007), the increase of H3K27ac in Hist2h3c2 is far higher than any other elevated Kac sites in the whole cell (e.g., H3K14ac increased 2.9-fold), showing a surprising 99.2-fold increase as compared to control (Fig. 1C). In addition, preliminary data from our lab characterizing H3K27ac in response to rotenone in the context of trained immunity indicated its significant alteration in a PD model. Accordingly, we speculated that H3K27ac plays an essential role in the cellular stress response of environmentally linked PD.

**Figure 1.**
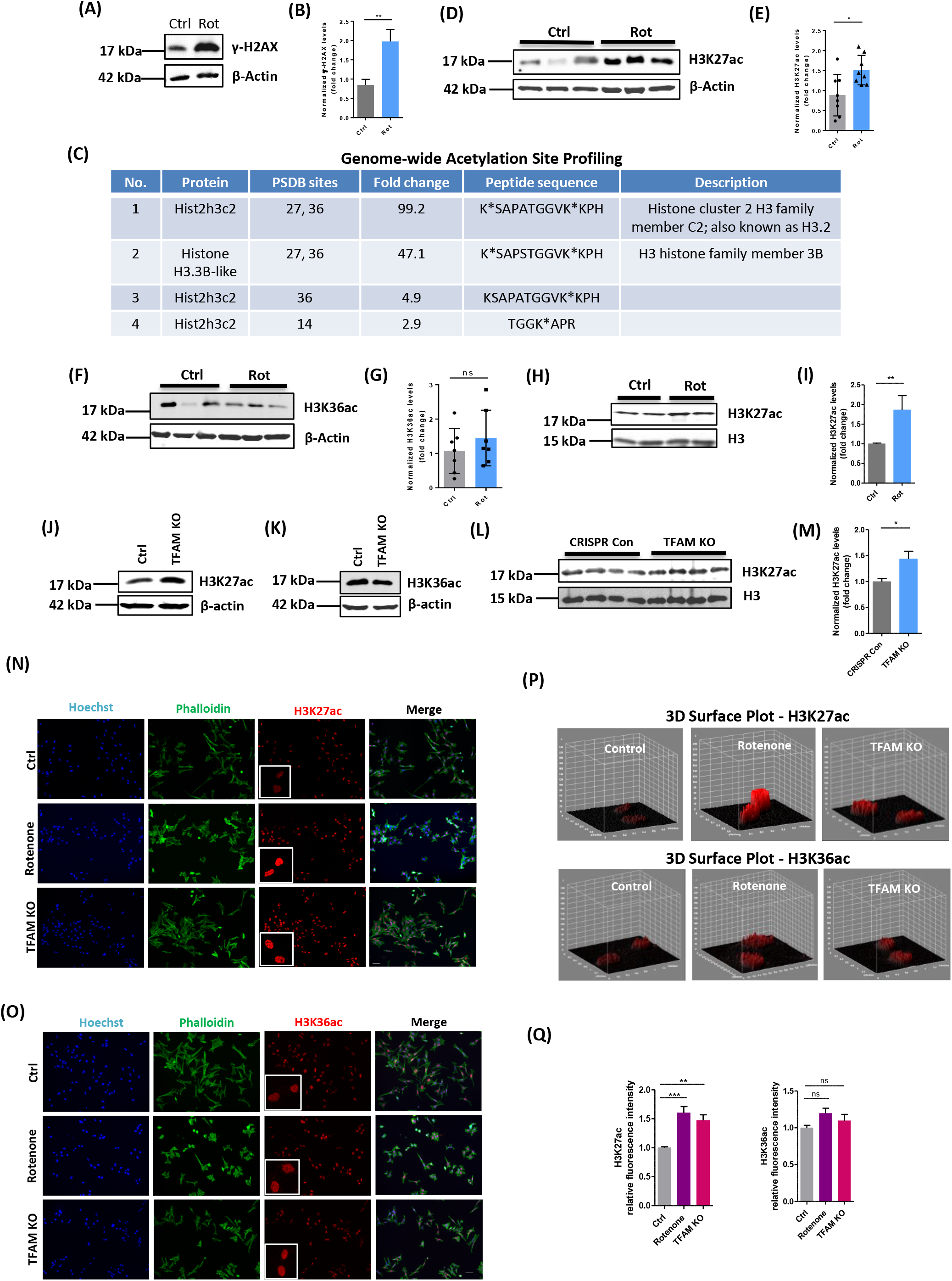
H3K27ac functions as the key acetylation site and responds to mitochondrial dysfunction in both rotenone-exposed and TFAM-KO DAergic neuronal cell models of PD. (A) Immunoblot validation of the expected DNA damage revealed by H2AX after 1 µM rotenone treatment for 3 h in AcetylScan samples; β-actin is the internal control. (B) Densitometric analysis of bands in immunoblot showing mean ± s.e.m of unpaired two-tailed t test (n=3). (C) Using LC/MS coupled with novel anti-acetyl-lysine antibodies and an optimized acetylome workflow for scanning the genome of rotenone-exposed N27 cells to map the acetylation sites. In the AcetylScan, only fold changes >2.5-fold were tracked with asterisks denoting acetylation sites (n=2). Peptide sequences of H3K27ac and H3K36ac are shown in histone cluster 2 H3 family member C2-encoded H3.2 and H3 family member 3B-encoded H3.3B. (D) Immunoblot and (E) quantification for H3K27ac from rotenone-treated N27 cells; β-actin is the internal control (n=8). (F) Immunoblot and (G) quantification for H3K36ac from rotenone-treated N27 cells. Five T175 flasks of cells were harvested for each sample. Independent experiments were repeated at least four times with n=7. Bar graphs show mean ± s.e.m of unpaired two-tailed t tests. (H) Immunoblots and quantification (I) of histone extracts from rotenone-treated N27 cells. Mann-Whitney test (n=4-6). (J) Immunoblot and (K) quantification for H3K27ac from TFAM-KO N27 cells. (L) Immunoblots and (M) quantification of histone extracts from TFAM-KO N27 cells. Unpaired t test followed by Welch’s correction (n=8). (N) Immunocytochemistry for H3K27ac and (O) H3K36ac from rotenone-treated and TFAM-KO N27 cells, with the nucleus stained with Hoechst (blue) and actin filaments with Phalloidin (green). Independent experiments were repeated at least four times. Scale bar, 50 µm. (P) 3D surface plot analysis of H3K27ac and H3K36ac in rotenone-treated and TFAM-KO N27s. (Q) Keyence BZ-X800 analysis of fluorescence intensity of H3K27ac and H3K36ac showing mean ± s.e.m of one-way ANOVA followed by Tukey’s post hoc test. ns, not significant; *p≤0.05; **p<0.01; ***p<0.001.

To eliminate the possibility that the high peak of the No.1 peptide was attributable to H3K36ac site rather than H3K27ac, we checked the peptide of a single H3K36ac site (Fig. 1 C). The single acetylation site (H3K36ac) had increased only 4.9-fold while the other H3K27ac peptide from histone variant H3.3B had increased 47.1-fold, indicating that H3K27ac is the highly induced Kac site responding to the mitochondria-impairing neurotoxicant and contributing to the corresponding gene expression reprofiling. Since H3K27ac and H3K36ac peaked together in both peptide sequences, it’s likely that H3K36ac facilitates H3K27ac to produce distinct outcomes in response to neurotoxicity, rather than H3K27ac alone reacting to cellular stress.

### H3K27ac mediates mitochondria-impairing cellular stress in neurotoxicant- exposed wild-type and TFAM-KO DAergic neuronal models of PD

To confirm the results of H3K27ac-mediated Kac site profiling in PD, we treated N27 cells with 1 μM rotenone for 3 h, and probed samples for H3K27ac and H3K36ac. The rabbit polyclonal antibodies that recognized the same epitopes within the internal region of H3K27ac and H3K36ac in AcetylScan were used. Immunoblot analysis revealed that H3K27ac increased significantly in rotenone-exposed, mitochondria-impaired DAergic neurons, while H3K36ac showed minimal accumulation (Fig. 1D-G). To further validate the H3K27ac accumulation, we extracted histones from the samples for immunoblots probed with H3K27ac (Fig. 1H-I), which were consistent with Fig. 1D. Therefore, we conclude that acetylation of the H3K27 site, and not that of H3K36, is the primary response to rotenone.

To further establish the elevated H3K27ac response to mitochondrial dysfunction, we added a distinct, complementary model for independent confirmation. TFAM (transcription factor A, mitochondrial) functions in genome transcription regulation and controls mitochondrial biogenesis (Miyazaki et al., 2012). Conditionally knocking out TFAM in DAergic cells produces mitochondrial dysfunction, which in mice leads progressively to a suite of neural and behavioral deficits that recapitulate PD (Langley et al., 2018). Using a CRISPR-Cas9 gene-editing method, we generated a stable TFAM-KO N27 cell line. With a >90% loss of TFAM protein confirmed with immunoblot (Fig. S2), H3K27ac was markedly elevated in TFAM-KO cells compared with CRISPR control cells (Fig. 1J). In addition, H3K36ac was not stimulated, as expected (Fig. 1K). Histone extracts of TFAM KO verifies this result (Fig. 1L-M).

Next, we used immunocytochemistry (ICC) analysis to characterize H3K27ac and H3K36ac in our rotenone and TFAM-KO models of PD. Both rotenone-exposed and TFAM-KO N27 cells displayed significantly increased signal intensities of nuclear H3K27ac (red in Fig. 1N, P-Q). However, the H3K36ac signal was much less inducible (Fig. 1O, P-Q), and therefore, much less responsible than was H3K27ac for the mitochondrial dysfunction. These data are consistent with our immunoblot results above.

To characterize mitochondrial impairment in rotenone and TFAM-KO models of PD, we used the mitochondrial membrane potential indicator JC-1, mitochondria-selective MitoTracker probes, Seahorse Bioscience’s extracellular flux analysis, and the mitochondrial superoxide indicator MitoSox. The JC-1 assay shows a decreased red/green ratio in the rotenone and TFAM-KO models (Fig. S3A-D), indicative of increased depolarization of the mitochondrial membrane. TFAM KO reveals higher cell number in green, indicating more neurons in early stages of cellular apoptosis (Fig. S3E). MitoTracker probe staining reveals increased mitochondrial circularity and damaged mitochondrial structure in the rotenone and TFAM-KO models (Fig. S4A-B). The functional consequences of the mitochondrial damage emerged as reductions in basal respiration rate, spare respiratory capacity, proton leak, and ATP production in the rotenone and TFAM-KO models as revealed by the extracellular flux analyses (Fig. S5A-C and S6A-C). A cell energy phenotype test shows that rotenone also stressed the oxygen consumption rate (OCR) and extracellular acidification rate (ECAR)/glycolysis (Fig. S5D), while TFAM KO averaged a lower OCR (Fig. S6D) relative to controls. Moreover, oxidative stress was induced in both rotenone-treated N27 and TFAM-KO N27 cells as indicated by the remarkably increased mitoROS production in the MitoSox assay (Fig. S7A-B). These data support the hypothesis that mitochondrial dysfunction characterizes both environmentally and genetically induced models of PD. We conclude that H3K27ac, rather than H3K36ac, acts as a critical player in mediating the cellular response to mitochondria-impairing stress and that mitochondrial dysfunction causes the hyperacetylation and the resulting impaired metabolism.

### Transcriptomic similarity of neurotoxicant-exposed and TFAM-KO DAergic neuronal cells identifies dysfunctional mitochondria in cellular stress response

Having established that mitochondrial dysfunction can lead to dramatic changes of chromatin structure, we next determined its consequence on the transcriptome. We performed RNA-seq analyses (three biological replicates) of rotenone-treated N27 and TFAM-KO N27 cells. Given that H3K27ac dramatically increases in both rotenone-treated N27 and TFAM-KO N27 cells, we indeed observed stunning numbers of genes either up- or down-regulated in both models (detailed below).

In our rotenone PD model, we observed that rotenone exposure upregulated 644 genes and downregulated 767 genes (Fig. 2A), based on a false discovery rate (FDR) cutoff of 0.05 (FDR ≤ 0.05) and a greater than 1.5-fold change (log_2_FC ≥ 0.585) in expression. We further validated our RNA-seq results with qRT-PCR analyses for five selected genes (i.e., *ADIPORT1, BTG2, GNAS, RNF2*, and *TMEM183a*) (Fig. 2B). These genes are related either to mitochondrial or neuronal functions. For example, deficiency of RNF2 increases autophagy induction (Xia et al., 2014). The imprinted *GNAS* gene can accelerate neuron death (Kim et al., 2006; Martos et al., 2017; Plagge et al., 2008). To understand the biological roles of the differentially expressed genes (DEGs) upon rotenone treatment, we performed Gene Ontology (GO) enrichment analysis for biological process. The most significantly enriched biological processes for these DEGs were mainly about regulation of the Wnt signaling pathway, NF-κB translocation, the cholesterol metabolic process, transcription regulation, protein function, and cell adhesion/differentiation (Fig. 2C).

**Figure 2.**
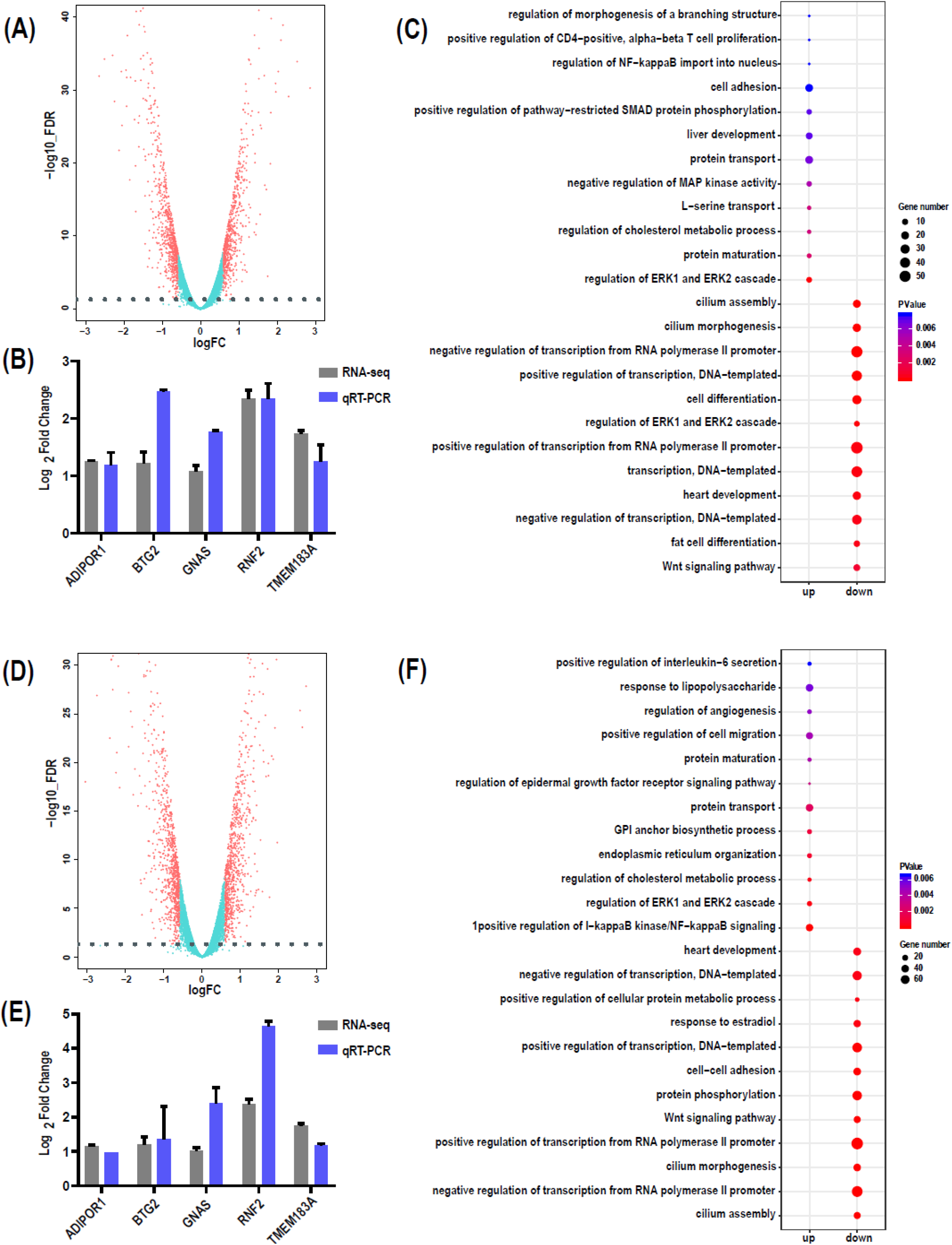
Transcriptomic alterations in rotenone-treated and TFAM-knockout (KO) N27 cells. (A) Volcano plot showing 644 genes upregulated (right) and 767 genes downregulated (left) upon rotenone treatment. (B) qRT-PCR validation of multiple differentially expressed genes (DEGs) upon rotenone treatment. (C) Gene Ontology analysis revealing the top 12 biological processes enriched by upregulated (left) and downregulated (right) genes upon rotenone treatment. (D) Volcano plot showing 813 genes upregulated (right) and 862 genes downregulated (left) by TFAM KO. (E) qRT-PCR validation of multiple DEGs upon TFAM KO. (F) Gene Ontology analysis revealing the top 12 biological processes enriched by TFAM KO.

In our TFAM-KO genetic model, we observed that TFAM KO upregulated 813 genes and downregulated 862 genes based on the same criteria applied to the rotenone group (Fig. 2D). We also validated our RNA-seq results from TFAM-KO N27 cells with the five genes selected above for qRT-PCR assay and confirmed their expression changes (Fig. 2E). GO enrichment analysis revealed that the most significantly enriched biological processes for these DEGs were mainly associated with protein maturation/transport, NF-κB and ERK1/2 signaling pathways, the cholesterol metabolic process, cilium structure, and IL-6 secretion (Fig. 2F).

With a large number of genes perturbed in both models (Fig. 2), we characterized the extent to which rotenone exposure and TFAM KO shared the effect on transcription. Data revealed that a stunning number of up- or down-regulated genes was shared in both models. The majority of upregulated genes (509 out of 645 rotenone- and 814 TFAM-KO-upregulated genes) were shared (Fig. 3A). Similarly, 619 genes were shared in 768 rotenone- and 862 TFAM-KO-downregulated genes (Fig. 3B). Further GO enrichment analysis showed that these overlapping downregulated genes are mainly enriched in the biological processes including ERK1/2 cascade regulation, NF-kB translocation and signaling, branching structure morphogenesis, and metabolic processes. Overlapping upregulated genes are associated with biological processes such as transcription regulation, cilium morphogenesis and assembly, and the Wnt signaling pathway (Fig. 3C).

**Figure 3.**
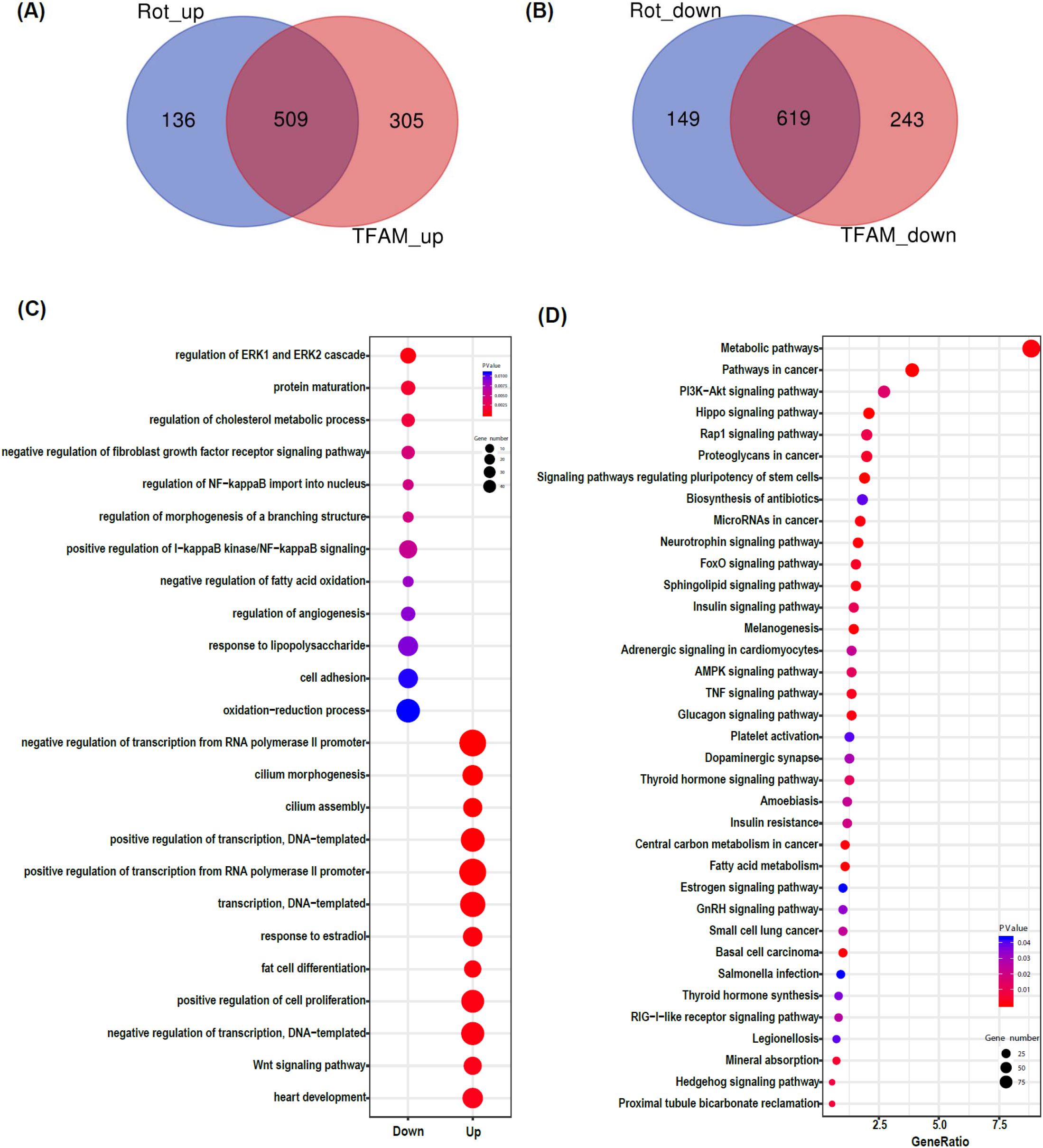
Common differentially expressed genes in rotenone-treated and TFAM-knockout (KO) N27 cells. Venn diagrams highlighting (A) upregulated and (B) downregulated DEGs shared between rotenone-treated and TFAM-KO N27s. (C) Gene Ontology analysis revealing the top 12 biological processes enriched by upregulated genes (left) and downregulated genes (right) overlapping between rotenone treatment and TFAM knockout. (D) KEGG pathway analysis identifying the most significant pathways for common, enriched DEGs.

In addition, Kyoto Encyclopedia of Genes and Genomes (KEGG) pathway annotation was performed to further explore the function of the parental genes of the DEGs and multiple pathways were identified (Fig. 3D). Intriguingly, the metabolic pathway is the most enriched pathway with many DEGs involved and the pathway of highest significance. Additional pathways highly relevant to PD include the apoptosis-related PI3K/AKT signaling pathway, the AMPK signaling pathway activated by falling energy levels yet exacerbating cell death (Curry et al., 2018), and the TNF signaling pathway associated with mediating neurotoxicity. Collectively, the enrichment further highlights our hypothesized axis of “exposure/mutations-mitochondrial dysfunction-metabolite-H3K27ac-transcriptome’ for PD pathogenesis.

### ChIP-seq identification of vulnerable genomic loci upon mitochondrial impairment

Our immunoblotting assays revealed globally increased levels of H3K27ac in mitochondria-impaired N27 cells. We next identified what epigenomic loci became more vulnerable to modification and what TFs were activated. We did ChIP-seq analyses in both rotenone-treated N27 and TFAM-KO N27 cells by using our previously characterized anti-H3K27ac (Wang et al., 2009). Genome-wide H3K27ac signals were analyzed following our established pipeline. We first determined the H3K27ac distribution at the genome level. Of the peaks identified for H3K27ac, approximately 25% intersected with annotated genes or their proximal promoters (hereafter defined as gene promoter regions located within 2.5 kb upstream and downstream of transcription start sites), though most peaks (75%) corresponded to intergenic regions (Fig. S8 A). The genomic distribution of H3K27ac did not differ significantly between the rotenone or TFAM-KO models.

When examining whether H3K27ac’s modification status was altered upon mitochondrial impairment, we found that 11.5% of H3K27ac peaks were significantly changed after rotenone treatment, and 8.0% of H3K27ac peaks were significantly changed after TFAM KO (Fig. S8 B). Regarding the genomic loci that were made more vulnerable in mitochondrially dysfunctional cells, 2.79% and 2.82% of H3K27ac peaks at annotated genes and their gene promoter regions were significantly changed after rotenone treatment and TFAM KO, respectively. Among these altered H3K27ac signals, we were particularly interested in genomic loci bound by the overexpressed histone mark H3K27ac. A total of 111 H3K27ac peaks located at gene promoter regions increased in rotenone-treated samples when compared to controls (Fig. 4A). Similarly, 88 H3K27ac peaks located at gene promoter regions increased in TFAM-KO samples when compared to controls (Fig. 4B).

**Figure 4.**
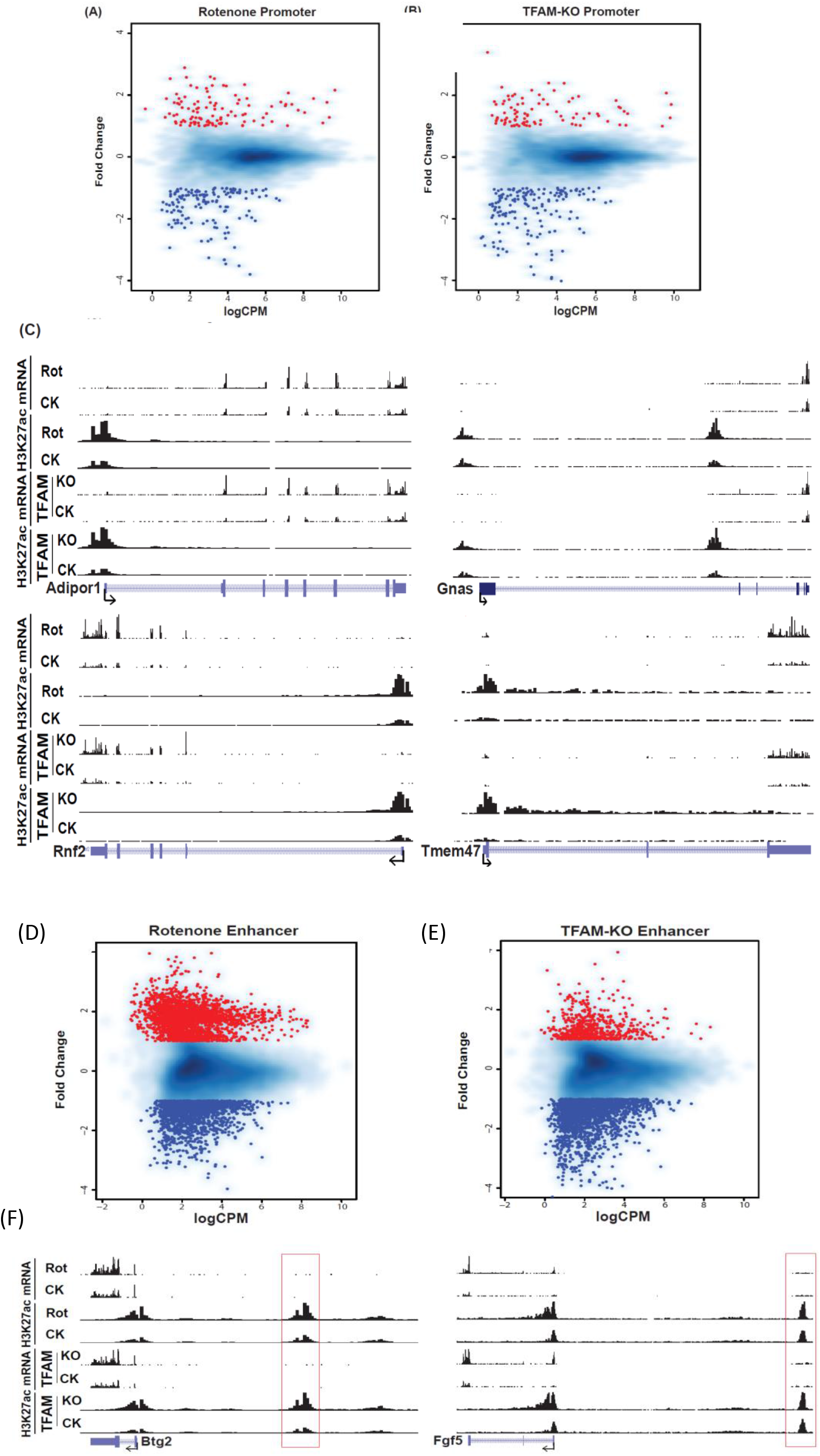
Identification of vulnerable genomic loci binding with H3K27ac and H3K27ac-specified enhancers in rotenone-treated and TFAM-knockout (KO) N27 cells. (A) Scatter plot showing H3K27ac peaks located at gene promoter regions in rotenone-treated and control N27 cell samples. (B) Scatter plot showing H3K27ac peaks located at gene promoter regions in TFAM-KO and CRISPR-control N27 cell samples. Each point represents a peak and the majority of points cluster in the middle. The dots indicate upregulated (red) and downregulated (blue) H3K27ac peaks. (C) Representative distribution of reads obtained by ChIP-seq and RNA-seq in four loci related to mitochondrial dysfunction and neural function in N27 cells. The distribution of H3K27ac was normalized to input and library dimension. (D) Scatter plot showing H3K27ac peaks located at enhancer regions in rotenone-treated and control N27 cell samples. (E) Scatter plot showing H3K27ac peaks located at enhancer regions in TFAM-KO and control N27 cell samples. (F) Representative distribution of reads obtained by ChIP-seq and RNA-seq in two gene loci and their distal promoter regions. Red rectangular boxes indicate H3K27ac-specified enhancers. The distribution of H3K27ac was normalized to input and library dimension. Uniform scales are used for H3K27ac and for mRNA. Rot: rotenone; CK: vehicle control; TFAM KO: TFAM knockout by CRISPR-Cas 9; TFAM CK: TFAM CRISPR control.

Having established that mitochondrial impairment increases H3K27ac, we next determined whether the H3K27ac modification was responsible for any gene expression changes, particularly increased expression (Wang et al., 2009; Wang et al., 2008). By integrating RNA-seq data, we found that an increase in H3K27ac was significantly associated with transcription promotion in N27 cells undergoing mitochondrial impairment. H3K27ac changes and consistent alteration of transcription across were shown at several genomic loci, illustrating mRNA expression levels sensitive to H3K27ac modification changes induced by our mitochondria-impairing models. We detected abundant H3K27ac in promoter regions together with upregulated mRNA levels at multiple gene loci such as *ADIPOR1*, *GNAS, RNF2, and TMEM47* (Fig. 4C). These data highlight the association between increased H3K27ac and elevated transcription of these genes.

### Characterization of genomic enhancers specific to mitochondria impairment and identification of corresponding transcription factor-binding motifs

Given that multiple genomic loci were influenced by mitochondrial impairment, we hypothesized that non-coding genetic regions like enhancers are affected as well. To identify such specific enhancers related to mitochondrial impairment, we further analyzed our ChIP-seq data using H3K27ac as an enhancer-specific histone modification mark.

After excluding H3K27ac peaks located at gene promoter regions, we found that 14.36% and 9.63% of H3K27ac peaks at distal promoters were significantly changed after rotenone treatment and TFAM KO, respectively (Fig. S8). We further identified 2336 and 601 promoter-distal and putative enhancer regions that increased in rotenone-treated N27 and TFAM-KO N27 cells, respectively (Fig. 4D-E). These results indicate that large numbers of enhancers defined by the H3K27ac modification mark were sensitive to mitochondrial impairment, which has implications for enhancer-specific gene regulation related to PD pathogenesis. Moreover, relatively more H3K27ac marked enhancers were affected by rotenone treatment than by TFAM KO, suggesting that the former is involved in more mitochondria-dependent pathways than the latter. Typical enhancers located upstream of *BTG2* and *FGF5* gene loci are presented in Fig 4F.

To identify TFs that interact with aberrant HAc caused by mitochondria impairment, we applied the HOMER tool to scan for TF-binding motifs within upregulated H3K27ac peaks at enhancer regions. We found that most of the enriched motifs in rotenone-upregulated H3K27ac peaks corresponded to TFs with a characteristic ZF (zinc finger-binding) domain or DM (zinc finger-like DNA-binding) domain (Fig. 5A). ZF proteins like ZNF264 contributed to the coordinated gene expression changes during brain aging (Hin et al., 2018) and ZNF711 is reportedly associated with mental retardation and cognitive disability (van der Werf et al., 2017). The DM family of TFs such as DMRT1 and DMRT3 were found to be involved in sexual development (Kopp, 2012). In contrast, bZIP motifs were more enriched in TFAM-KO-upregulated H3K27ac peaks (Fig. 5B). The family of bZIP TFs has been assigned important roles in cancer development, hormone synthesis, and reproductive functions (Hoare et al., 1999; Manna et al., 2002; Vlahopoulos et al., 2008). In particular, AP-1, composed of heterodimers of Fos and Jun, activates SNpc microglia neuroinflammation and drives DAergic neurons toward apoptosis (Tiwari and Pal, 2017).

**Figure 5.**
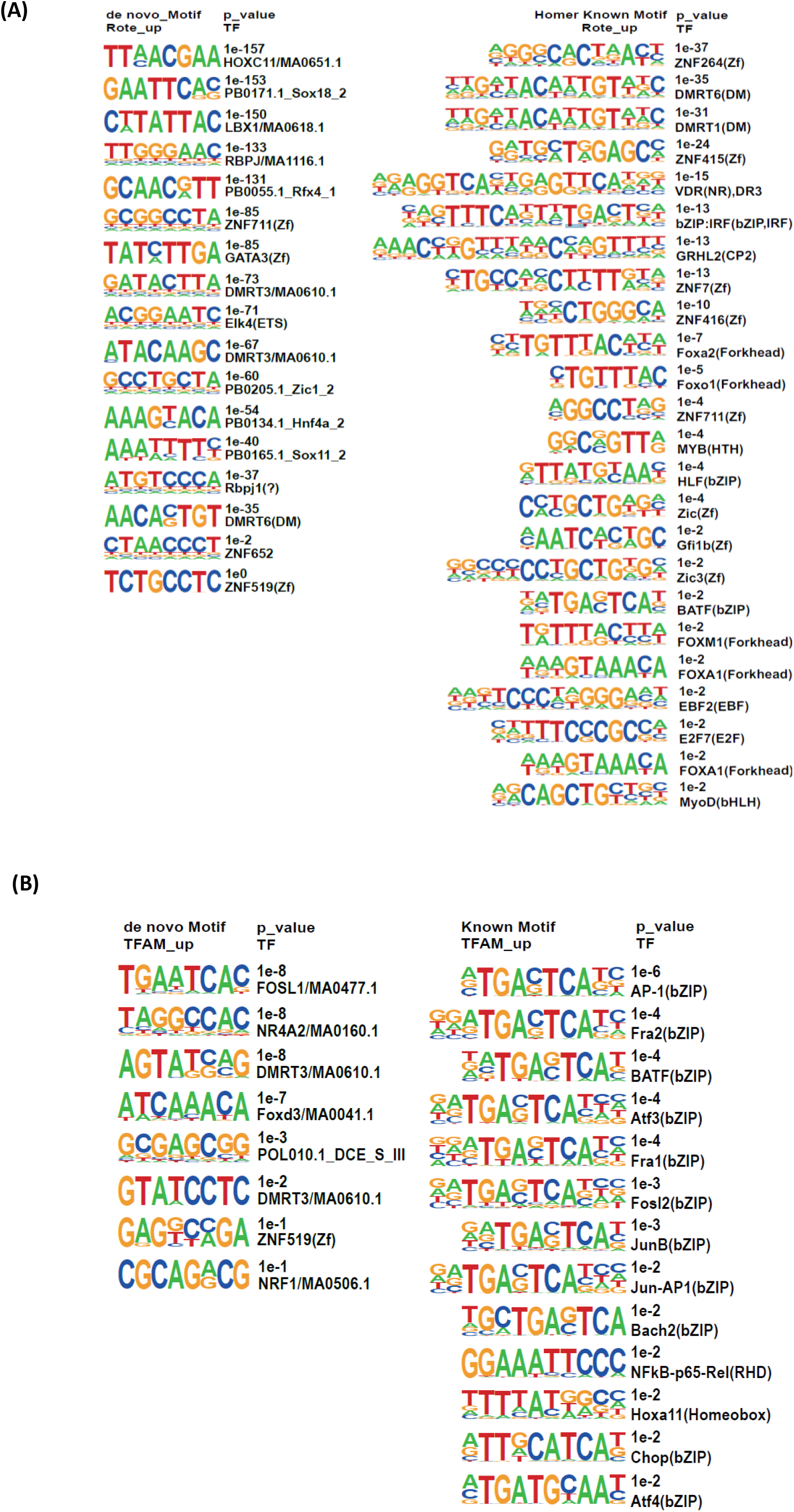
Motif analysis in rotenone-exposed and TFAM-KO N27 cells displaying motifs at (A) rotenone-affected enhancers or (B) TFAM-KO-affected enhancers. The top enriched *de novo* sequence motifs appear in the left panel with their best-known motif match names, while the top enriched known motifs appear in the right panel.

To further explore the unknown motif landscape of enhancers, we applied *de novo* motif discovery on upregulated enhancers related to rotenone exposure and TFAM KO (Fig 6A and B). Although these novel motifs were lacking perfect database matches, HOMER provided similar known motifs which most closely matched the *de novo* motifs. The most enriched motif in rotenone-upregulated enhancers was related to the homeobox family, playing an important role in morphogenesis in all multicellular organisms (Zhang et al., 2007). The other enriched motif in rotenone-upregulated enhancers was closely matched with the RBPJ (recombination signal-binding protein for immunoglobulin kappa J region) TF, which was reported to control the Notch signaling pathway by recruiting histone deacetylase or histone acetyltransferase (Borggrefe and Oswald, 2009). The most enriched motif in TFAM-KO-upregulated enhancers was matched with FOLS1 (Fos like 1), a component of AP-1 TF complexes and contributes to the regulation of placental development (Kubota et al., 2015). By revealing the known and unknown TF motifs in neurodegeneration driven by mitochondrial dysfunction, our investigations have shown the intricate connections between epigenetic and transcriptional regulation, providing unprecedented insights into understanding PD pathogenesis.

**Figure 6.**
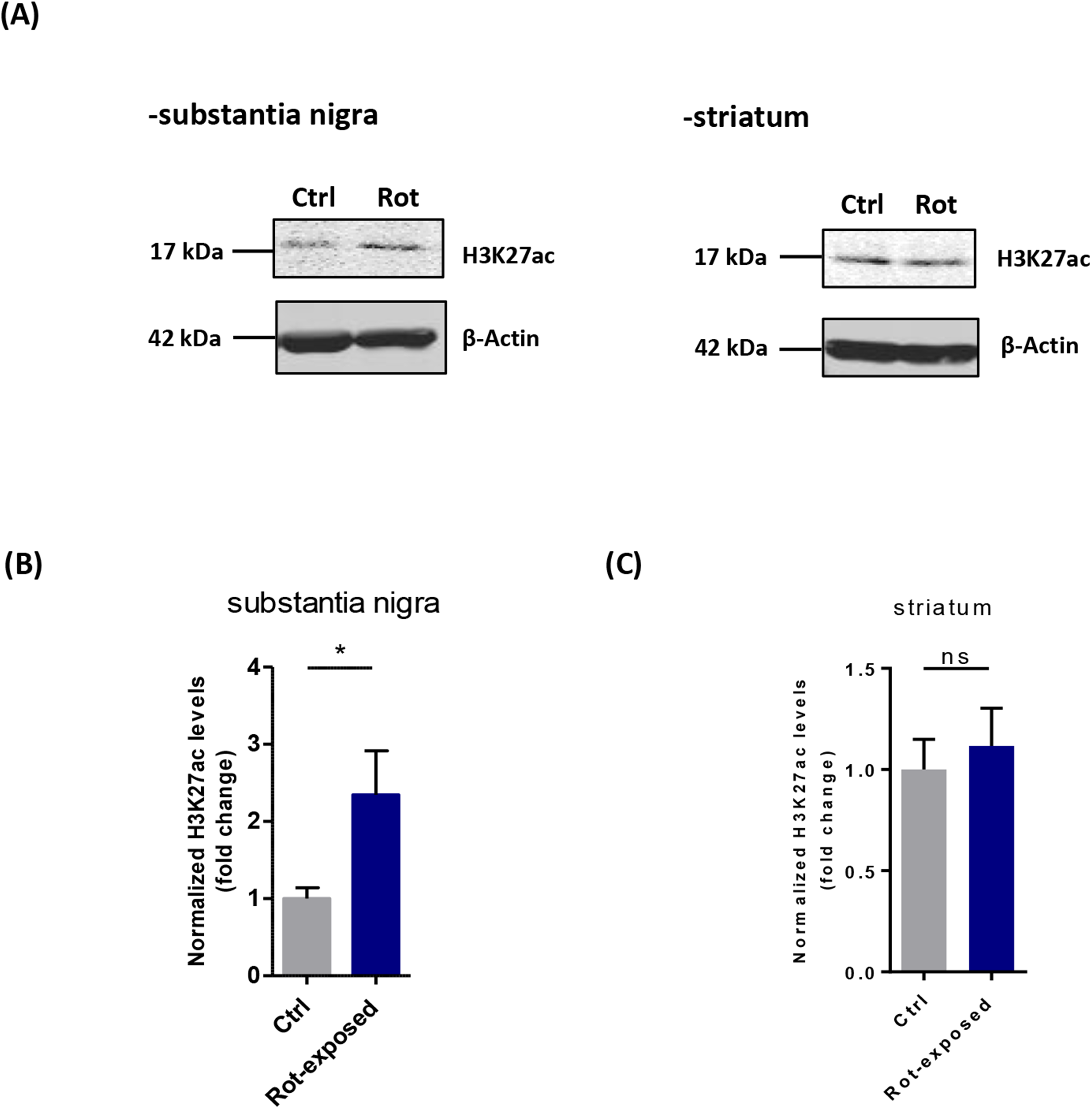
Elevated H3K27ac in substantia nigra (SN) of MitoPark mice and rotenone-exposed ex-vivo midbrain slices. (A) Representative immunoblots for H3K27ac in rotenone-exposed midbrain slices coupled with their densitometric analyses for the (B) SN and (C) striatum. Each data point is the average of three replicates. At least four independent experiments were measured for SN and two for striatum. Independent experiments were repeated four times. Bar graphs show mean ± s.e.m of unpaired two-tailed t tests. ns, not significant; *p≤0.05; **p<0.01.

### Mitochondrial dysfunction induces H3K27ac to mediate mitonuclear communication and the loss of DAergic midbrain neurons

The bidirectional regulation between mitochondria and the nucleus, defined as mitonuclear communication, maintains cellular homeostasis and cellular adaptation to environmental stress. Retrograde response signaling from mitochondria to the nucleus adjusts gene expression through epigenomic modifications (Matilainen et al., 2017; Quirós et al., 2016). Having shown that mitochondrial dysfunction induced H3K27ac to mediate retrograde responses and subsequently alter gene expression *in vitro*, we then extended these findings *ex vivo* to rotenone-exposed acute midbrain slices as well as *in vivo* to the transgenic MitoPark mouse model. The analysis demonstrated mitochondrial dysfunction in treated slices by using COXIII and TFAM as biomarkers (Fig. S9A). Immunoblot shows significant increase of H3K27ac in the SN, but not the striatum. This was consistent with our *in vitro* findings that mitochondria-impairing stress triggers mitonuclear signaling that induces H3K27ac (Fig. 6A-C).

To examine the impact of H3K27ac on gene expression *ex vivo*, several upregulated candidate genes were selected from our integrated analyses of RNA-seq and ChIP-seq. Our qPCR analyses of the SN region confirmed notably enhanced mRNA levels of *TMEM47* in midbrain slices (Fig. S9B), which is in accordance with our H3K27ac reads at genomic loci for neuronal models (Fig. 4C and F). As a transmembrane protein, *TMEM47* is reported to negatively regulate Fyn kinase (Dong and Simske, 2016). Our previous publication shows that Fyn kinase, along with the class B scavenger receptor CD36, regulates the uptake of aggregated human α-synuclein (αSyn), the abnormal protein deposits in PD-associated Lewy bodies (Panicker et al., 2019). However, the specific function of *TMEM47* in human PD is not clear. *BTG2* also reveals increased transcription levels. *BTG2* targets p53 that negatively regulates cell cycle progression in response to DNA damage and other stressors, and is involved in mitochondrial depolarization (Fig. S3) (Ficazzola et al., 2001).

## Discussion

The integrated genome-wide analysis of H3K27ac distribution and transcriptomic alteration shed light on our hypothesis that H3K27ac plays a critical role during environmentally induced PD pathogenesis. It provides a novel paradigm for studying the epigenetic effects of chronic neurotoxicant exposure in neurondegeneration and reveals H3K27ac as a key epigenetic post-translational modification marker in environmentally linked PD. Altogether, our results reveal a pivotal axis of ‘exposure/mutation-mitochondrial dysfunction-metabolism-H3K27ac-transcriptome’ for PD pathogenesis (Fig. 7).

**Figure 7.**
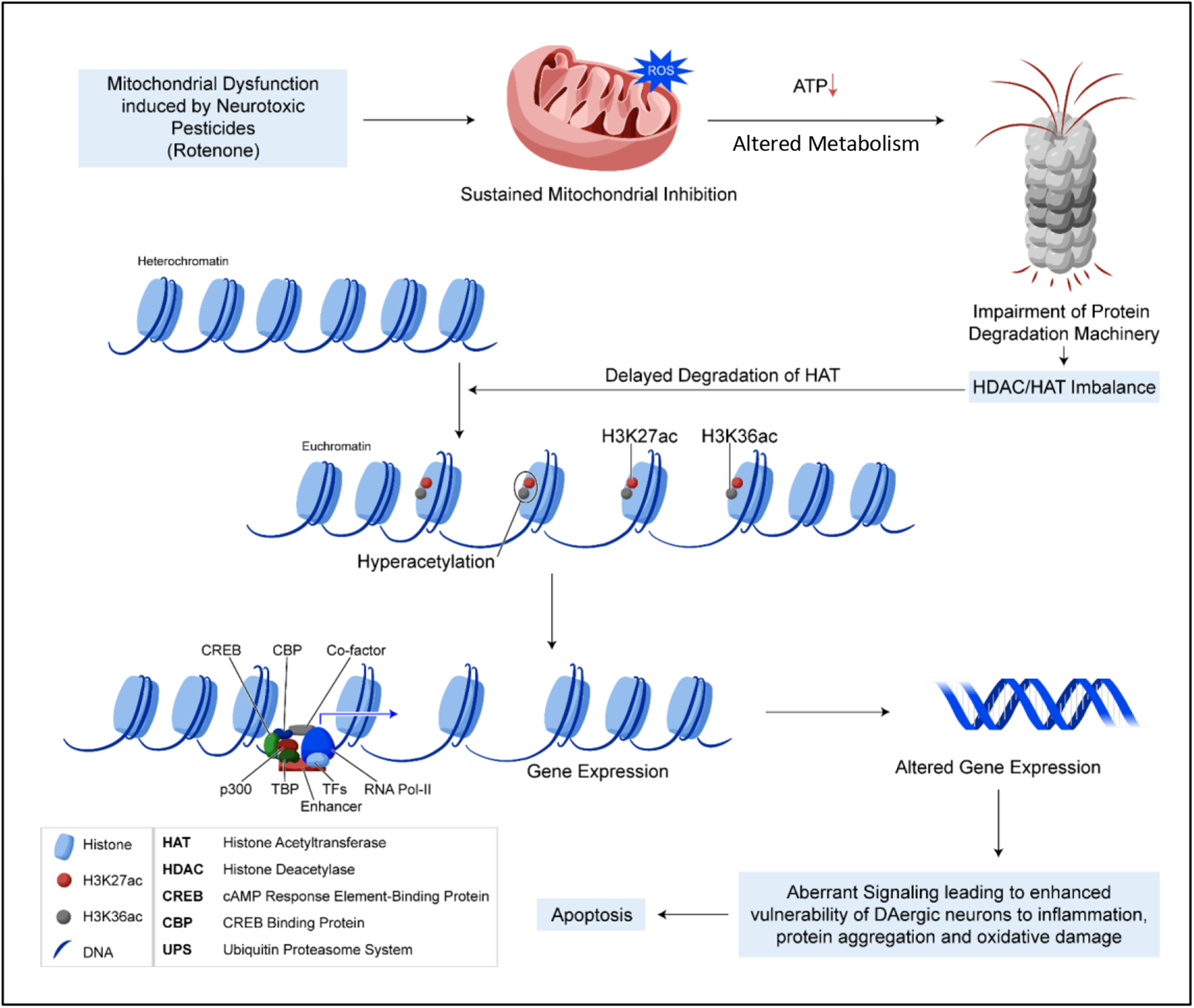
Mitochondrial dysfunction induces epigenetic dysregulation by H3K27 hyperacetylation to perturb active enhancers in Parkinson’s disease models

To the best of our knowledge, ours is the first study to integrate RNA-Seq and ChIP-Seq data for an in-depth analysis of genomic regions with differential chromatin activity in mitochondria-impairing PD models involving rotenone-treated N27 and TFAM-KO N27 neuronal cells. With this combined approach, we were able to confirm H3K27ac as a key factor involved in the epigenetic regulation of gene expression in mitochondria-impaired N27 cells. Through a series of biochemical studies, epigenomic profiling, and bioinformatic identification, our results reveal an axis of exposure and/or genetic mutation-mitochondrial dysfunction-metabolism-H3K27ac/active enhancer activity-transcriptome underlying the neurodegeneration of PD. In our opinion, this working model uncovers a mechanism of mitochondrial dysfunction common to both environmental neurotoxicant- and genetic mutation-driven PD. The latter includes genetic mutations of Parkin and PINK1. Our results were consistent at the chromatin structure and transcriptome levels using both the rotenone exposure and TFAM-KO models of PD, thus providing a robust test for the working model. On the other hand, the exposure model seemed to involve more complex pathways than the genetic model.

Comparing rotenone-treated and TFAM-KO N27 neuronal cell models of PD further revealed that an environmental neurotoxicant and a genetic mutation can induce similar transcriptomic changes that converge on shared gene expression changes associated with specific functions. We found evidence for shared functional processes and pathways among DEGs in these two models. Among upregulated genes, strong enrichment occurred in biological processes related to Wnt signaling pathway, transcription regulation, cilia functions, and estradiol responses, all of which have previously been shown to be associated with PD dysfunctions (Gonzalez-Reyes et al., 2012; Siani et al., 2017; Stephano et al., 2018). For example, impaired Wnt signaling in DAergic neurons has been associated with pathogenesis in a rotenone-triggered PD model (Stephano et al., 2018). Loss of cilia will decrease the ability of cholinergic neurons to respond to a sonic hedgehog signal that triggers neuroprotective signaling toward DAergic circuits (Gonzalez-Reyes et al., 2012). Downregulated genes also converged on shared functional categories, such as ERK1/2 regulation, protein maturation, cholesterol metabolic processes, and NF-κB translocation.

It is possible that rotenone neurotoxicity was largely exerted on DAergic neurons through impaired mitochondrial functions. To examine the epigenomic aberrations in mitochondria-impairing PD models, genome-wide H3K27ac profiling by ChIP-Seq was analyzed for differences between rotenone treatment and vehicle control or between TFAM KO and CRISPR control. Overall, specific H3K27ac signatures were identified at promoters and enhancers acting as regulatory elements associated with mitochondrial impairment. Among the identified differential H3K27ac peaks, <3% were located at gene promoter regions in either rotenone-exposed N27 cells or TFAM-KO N27 cells. Several genomic loci appeared to be consistent alteration targets of promoter-H3K27ac modification and gene expression in both model systems tested. *RNF2* gene expression was reported to be dysregulated in striatal lesioned brain hemispheres (Sodersten et al., 2014). *TMEM47* has been linked to neuronal development and/or brain tumors (Aruga et al., 2003; Utami et al., 2014). This appears to be the first report that these genes are affected in mitochondria-impairing PD models, as well as the corresponding H3K27ac modifications on these gene loci. It is likely that the mRNA expression levels of these gene loci are sensitive to promoter-H3K27ac modification changes resulting from mitochondrial impairment.

In addition to promoter-H3K27ac aberrations, we observed relatively more differential H3K27ac peaks at intergenic regions upon mitochondrial impairment. This is consistent with previous studies. H3K27ac is considered to be the representative acetylation mark to highlight active enhancers (Creyghton et al., 2010) and is more predictive of enhancers than other histone marks (e.g., H3K4me1, H3K4me2, H3K4me3 and H3K9ac) (Fu et al., 2018). Enhancers are non-directional regulatory elements containing TF-binding motifs. Upon binding with TFs, complexes are formed and alter the 3D conformation of chromatin to promote the transcription of target genes located in *cis* (Kumar et al., 2013). Population-scale genetic studies indicate that many sequence variants associated with human diseases reside in enhancers (Ernst et al., 2011; Maurano et al., 2012). However, little is known about H3K27ac-defined enhancers and their activity patterns in PD. Therefore, we globally interrogated the H3K27ac histone acetylome of enhancers in mitochondria-impairing PD models. We found that large numbers of H3K27ac-defined enhancers were highly sensitive to the mitochondrial dysfunction resulting from rotenone exposure or TFAM KO. Among the TFs highlighted by motif analysis of upregulated H3K27ac peaks, the ZF factors ZNF264 and ZNF711 show a strong genetic association with neurological functions (Hin et al., 2018; van der Werf et al., 2017). AP-1, another enriched TF, reportedly promotes microglial neuroinflammation and DAergic neuron apoptosis in the SNpc (Tiwari and Pal, 2017). Interestingly, AP-1 has also been identified as a key TF enriched by H3K27ac in the brain tissues of a large cohort of autism spectrum disorder patients (Sun et al., 2016). Furthermore, apart from these known motifs, our *de novo* motif discovery identified novel non-canonical motifs, providing additional insight into TF-binding preferences. These results suggest that mitochondrial dysfunction, whether induced through neurotoxicant exposure or genetic mutation, influences the binding of distal H3K27ac defined enhancers (regulatory elements), thus affecting the gene expression contributing to the etiology of PD. Future investigations untangling metabolomic changes may prove equally insightful in charactering elevated H3K27ac in mitochondria-impairing PD model systems.

## Methods

### Chemicals and reagents

RPMI, neurobasal medium, fetal bovine serum (FBS), L-glutamine, IR-dye tagged secondary antibodies, Hoechst nuclear stain, penicillin, streptomycin, horse serum, and other cell culture reagents were purchased from Invitrogen (Gaithersburg, MD). The Bradford protein assay kit was purchased from Millipore. The Phalloidin antibody was ordered from Thermo Fisher Scientific, Protease and Phosphatase inhibitor cocktail from Life Technologies, Acetylation inhibitor cocktail from Santa Cruz, and propidium iodide (PI) from Molecular Probes.

### AcetylScan proteomic profiling

AcetylScan proteomic profiling followed the standard protocol as described by Svinkina et al. (2015). At least 10 T175 flasks of N27 DAergic neuronal cells (seeding density: 6×10^6^) were grown for each sample (n=2) with 2×10^8^ of N27 cells being cultured as one sample. Two samples were used as control. Another two samples of the same cell passage were collected and exposed to treatment. After growing to 70% confluency, cell viability and sub-confluence were inspected by microscopy. Next, cells were treated with 1 uM rotenone for 3 h. After treatment, cell harvesting proceeded until 10 flasks of cells had been collected. Adherent cell cultures were washed by adding 10 mL of 4°C PBS. After pipetting off all PBS, 10 mL of urea lysis buffer (ULB: 20 mM HEPE pH 8.0, 9.0 M Pierce Sequanal grade urea (Cat. No. #29700), 1 mM sodium orthovanadate (activated), 2.5 mM sodium pyrophosphage, and 1 mM beta-glycerol-phosphate) was added to flask#1 before scraping off the cells. For the remaining 9 flasks, steps were the same except the ULB came from the previous flask (e.g., flask #2 used ULB containing proteins from flask#1).Protein was digested and desalted using pre-conditioned C18 solid phase extraction. Peptides were then eluted from antibody beads with 50 µL of 0.15% (TFA), and completely dried by lyophilization. Next, peptides were resuspended and enriched by motif anti-Kac antibodies noncovalently coupled to protein A agarose beads and triple-washed with 1.5 mL buffer (Cat. No. #13416). After immunoprecipitation of beads, the resin was washed and peptides were eluted from it. Peptide fractions were analyzed by online nanoflow liquid chromatography tandem mass spectrometry (LC-MS/MS). MS parameter settings were as follows: MS Run Time 96 min, MS1 Scan Range (300.0 – 1500.00), Top 20 MS/MS (Min Signal 500, Isolation Width 2.0, Normalized Coll. Energy 35.0, Activation-Q 0.250, Activation Time 20.0, Lock Mass 371.101237, Charge State Rejection Enabled, Charge State 1+ Rejected, Dynamic Exclusion Enabled, Repeat Count 1, Repeat Duration 35.0, Exclusion List Size 500, Exclusion Duration 40.0, Exclusion Mass Width Relative to Mass, Exclusion Mass Width 10 ppm). For data analysis, sequences were assigned to MS/MS spectra with Sorcerer for relative quantitation. MS/MS spectra were evaluated using SEQUEST (J.K Eng, 1994, J Am Soc Mass Spectrom) and the core platform from Cell Signaling. Searches were performed against the most recent update of the NCBI database with mass accuracy of +/-50 ppm for precursor ions and 1 Da for productions. Results were filtered with mass accuracy of +/– 5 ppm on precursor ions and presence of the intended motif. Data were presented by fold change in protein types, which used a 2.5-fold cutoff to determine significant change.

### MTS mitochondrial activity assay

At 1 h, 3 h and 6 h post-treatment, 10 μL of MTS dye (Promega) was added to cells in 96-well plates and incubated for 1 h. After incubation, the plates were read at 490 nm (excitation) and 680 nm (background), respectively.

### CRISPR/Cas9-based TFAM-KO cell generation

The lentivirus-based CRIPSR/Cas9 TFAM-KO method follows Gordon et al. (2016). The TFAM-KO plasmid pLV-U6gRNA-Ef1aPuroCas9GFP-TFAM, with the TFAM-gRNA target sequence directed against the exon 1 sequence (CPR555e5e4099bf84.98), was purchased from Sigma-Aldrich. The lenti-CRISPR/Cas9 TFAM-KO plasmid and universal negative control lentivirus (U6-gRNA/CMV-Cas9-GFP) were respectively transfected into 293FT cells using the Mission Lentiviral Packaging Mix (Cat#SHP001, Sigma-Aldrich) according to manufacturer’s instructions. The lentivirus was harvested 48 h after transfection and added to N27 cells for infection at an MOI of 100. After 24 h, puromycin (50 μg/mL) was supplemented for stable cell selection.

### Mitochondrial membrane potential, morphology, and superoxide production

JC-1 mitochondrial potential sensor was purchased from Invitrogen and the standard commercial protocol was followed. Briefly, 2.0 μg/mL of JC-1 diluted in serum-free N27 media was added to N27 cell cultures and incubated at 37°C for 20 min. Following gentle, triple washes with PBS, images were immediately captured by Keyence microscope before cells dried out. The ratio of red to green was calculated. For MitoTracker staining, procedures were similar. Following treatment paradigm, 300 μl of 166-nM CMXROS MitoTracker red dye diluted in serum-free N27 media was added and incubated at 37°C for 12 min. After triple-washing with PBS, cells were fixed in 4% PFA for 30 min prior to following the steps for ICC and 3D imaging. For MitoSox, live N27 cells were stained following the manufacturer’s instructions to detect superoxide production.

### Mitochondrial bioenergetics analysis

To measure mitochondrial oxygen consumption and extracellular acidification rates, a Seahorse XFe24 analyzer was used for the Mito Stress test. Following Panicker et al. (2019), 0.75 μM oligomycin, 1 μM carbonyl cyanide p-trifluoromethoxyphenylhydrazone, and 0.5 μM rotenone-antimycin, serum-free medium were used.

### Animal studies

All animal studies followed animal procedures approved by Iowa State University’s (ISU, Ames, IA, USA) Institutional Animal Care and Use Committee (IACUC). All mice used for this study were bred and maintained under a 12-h light cycle in a climate-controlled mouse facility (22±1°C) with food and water available *ad libitum*.

### Acute midbrain slice

Use of the animals and protocol procedures were approved by the IACUC at ISU. Organotypic slices were prepared as previously described with several steps improved to adapt to a PD disease model (Kondru et al., 2017). Slice culture buffer Gey’s balanced salt solution supplemented with the excitotoxic antagonist kynurenic acid (GBSS) was prepare as the following order: 8g NaCl, 0.37g KCl, 0.12g Na_2_HPO_4_, 0.22g CaCl_2_∙2H_2_O, 0.09g KH_2_PO_4_, 0.07g MgSO_4_∙7H_2_O, 0.210 g MgCl_2_∙6H2O, 0.227g NaHCO_3_. WT (C57BL/6) postnatal mouse pups (7-to 12-day old) were used as source for live brain slices. The microtome’s (Compresstome VF-300, Precisionary Instruments) slicing reservoir was prefilled with freshly prepared, ice-cold sterile GBSS. Dissected midbrain from the whole brain was oriented in the sagittal plane in the Compresstome’s specimen tube with 2% liquid agarose inside. After quickly solidifying the agar by clasping the specimen tube with a chilling block, the specimen tube was inserted into the slicing reservoir. Nigrostriatal slices (200 μm) were transferred to 6-well plates with oxygenated GBSS. The plates were preheated to 37°C in a humidified atmosphere of 5% CO2. After a 1-to 2-h recovery, acute slices were treated with 25 µM rotenone for 3 h. Afterward, slices were gently double-washed in ice-cold PBS, temporarily stained with propidium iodide to ensure viability and homogenized to prepare lysates for Western blotting or RNA for qRT-PCR.

### SYBR Green qRT-PCR

Slices were exposed to rotenone treatment in 6-well plates in an incubator maintained at 5% CO2 and 37°C. Treatments were performed in serum-free RPMI media supplemented with penicillin (100 U/ml), streptomycin (100 ug/ml), and 2 mM L-glutamine. After a 48-h treatment, slices were washed with ice-cold PBS and harvested. RNA was isolated from slices according to Seo et al. (2014). In brief, slices were lysed in 1 mL TRIzol reagent and incubated for 5 min at room temperature, then 0.2 mL chloroform was added to tubes and incubated for 2 min. Next, tubes were centrifuged at 12,000 x g for 15 min at 4°C. The top layer in the tubes containing RNA were transferred to a new tube containing 0.7 mL isopropanol. After a 15-min incubation, tubes were again centrifuged at 12,000 x g for 10 min at 4°C to precipitate RNA. Later the supernatant was discarded, and RNA pellets were washed with 75% ethanol. After air-drying to completely remove any residual ethanol, RNA pellets were dissolved into water. NanoDrop was used to test their concentration, and 1000 ng of RNA was utilized to convert into cDNA. The high capacity cDNA synthesis kit from Applied Biosystems (#4368814) was used for cDNA synthesis according to manufacturer’s protocol. For normalization of each sample in qRT-PCR, the 18S rRNA gene (Cat. No. PPM57735E) was used as the housekeeping gene. Based on manufacturer’s guidelines, dissociation curves and melting curves were run to ensure that single amplicon peaks were obtained without any non-specific amplicons. Fold change in gene expression was determined using the ΔΔC_t_ threshold cycle (C_t_) method. Quantitative PCR was performed using qRT^2^PCR SYBR Green Mastermix (Agilent). Primers were ordered from Qiagen.

### Histone extraction

Histone extraction was performed according to Song et al. (2010). In brief, samples were incubated with cytosolic extraction reagent I provided by a NE-PER kit (Thermo Fisher Scientific) and 0.5% Triton X-100 for 10 min according to the commercial protocol. Tissue homogenization was processed before incubation. After incubation, the pelletized nuclei were collected following centrifugation at 2000 x g for 5 min and then resuspended in 0.2 N HCl. Overnight incubation at 4°C was performed for complete dissolution of histones. The following morning, samples were centrifuged at maximum speed for 10 min and then the pellets were collected.

### Immunoblotting

After rotenone treatment, cells were lysed using modified RIPA buffer, homogenized, sonicated, and centrifuged according to Kaul et al. (2003). For treated tissue, slices were lysed with modified RIPA, homogenized, and centrifuged. Proteins were normalized using Bradford assay before loading on Sodium Dodecyl Sulfate (SDS)-acrylamide gels. For separation of proteins, 20-40 μg of protein was loaded in each well of 15% acrylamide gels. Acrylamide gels were run at 103 V for 2 .5 h at 4°C. Following separation, proteins were transferred to a nitrocellulose membrane at 25 V overnight at 4°C. After transfer, the membranes were blocked with LI-COR blocking buffer for 1 h and incubated in primary antibodies following manufacturer’s protocol. Next, membranes were washed with PBS-TWEEN 20 (0.05%) for 1 h and incubated in LI-COR IR secondary antibodies for 1 h on a belly dancer at room temperature. After washing with PBS-TWEEN for 1 h, membranes were imaged using an Odyssey scanner. The primary antibodies used are as follows: H3K27ac (Cell Signaling, 1:1000), gamma-H2AX (Cell Signaling, 1:1000), TFAM (Santa Cruz, 1:1000), H3 (Millipore, 1:1000) and β-actin (Sigma, 1:10000) as loading control. The secondary antibodies used are as follows: IR-800 conjugated goat anti-mouse IgG (LI-COR, 1:20000), IR-700 conjugated goat anti-rabbit IgG (LI-COR 1:20000), and IR-800 conjugated donkey anti-goat IgG (LI-COR, 1:20000).

### ICC

For ICC, 4% PFA was used to fix N27 DAergic neuronal cells. After a double-wash, fixed neurons were blocked using blocking buffer, and incubated in primary antibodies following manufacturer’s protocol. Following primary antibody (H3K27ac 1:500, Cell Signaling) incubation, cells were washed with PBS and incubated in Alexa dye-conjugated secondary antibody. Next, neurons were washed and mounted on slides using Fluoromount aqueous mounting medium (Sigma). Neurons were visualized using an inverted fluorescence microscope (Nikon TE-2000U).

### RNA-seq library construction and sequencing

Total RNAs from treatment and control groups were isolated and quality controlled according to the Illumina protocols. A total of 2 μg RNA per sample was used as initial material for library construction. Poly-A containing mRNA molecules was purified by using poly-T oligo-attached magnetic beads and fragmented into small pieces using divalent cations under elevated temperature. The cleaved RNA fragments are copied into first strand cDNA using reverse transcriptase and random primers. Strand specificity is achieved by replacing dTTP with dUTP, followed by second strand cDNA synthesis. These cDNA fragments then have the addition of a single ‘A’ base and subsequent ligation of the adapter. The products are then purified and enriched with PCR to create the final cDNA library. RNA-seq was performed on an Illumina Hiseq X10 platform and 150-bp paired-end reads were generated according to Illumina’s protocol. RNA-seq data were submitted to the GEO database.

### Whole-genome gene expression analysis

Data analysis was performed similar to Li et al. (2015). The adapter sequences were removed from the raw sequencing data and the individual libraries were converted to the FASTQ format. Sequence reads were aligned to the rat genome (rn6) with HISAT2 (v2.1.0) and the resulting alignment files were reconstructed with the EdgeR3 package. RefSeq database (Build 37.3) was chosen as the annotation references for mRNA analysis. The read counts of each transcript were normalized to the length of the individual transcript and to the total mapped fragment counts in each sample and expressed as reads per kilobase of exon per million fragments mapped (RPKM) of mRNAs in each sample. The mRNA differential expression analysis was applied to treatment and control groups. An adjusted *p* value ≤ 0.05 (Student’s *t*-test with Benjamini-Hochberg FDR adjustment) was used as the cut-off for significantly differentially expressed genes. DEGs were analyzed by enrichment analysis to detect over-represented functional terms present in the genomic background. GO and KEGG pathway analysis were performed using the DAVID Bioinformatics Resources 6.8 (Huang da et al., 2009; Kanehisa and Goto, 2000; Valentine et al., 2019).

### Gene expression validation by quantitative reverse transcription-PCR (qRT-PCR)

Total RNAs were isolated and reverse transcribed into cDNAs by using PrimeScript RT reagent kit with gDNA Eraser (Takara Bio Inc., Japan). Five genes related to mitochondria and neurological functions were selected for qRT-PCR assay. Primers are listed in Table 1. All qPCR reactions were performed on an Applied Biosystems StepOne Real-time PCR system (Thermo Fisher Scientific, USA) using iTaq Universal SYBR Green Supermix (Bio-Rad, USA), with three technical replicates. The amplification procedure was as follows: 95°C for 5 min, followed by 40 cycles of 95°C for 10 sec and 60°C for 20 sec. Relative quantification of target genes was performed using the ΔΔC_t_ method with GAPDH as a reference gene.

### Chromatin immunoprecipitation plus deep sequencing (ChIP-seq)

ChIP was performed according to Wang et al. (2009). In brief, cells were cross-linked with 1% formaldehyde for 10 min at room temperature and quenched with 125 mM glycine for 10 min. After fragmenting into 100 – 1000 bp by sonication, chromatin templates from 3 million cells were used for ChIP assay. Samples were immunoprecipitated with 2 μg H3K27ac antibody overnight at 4°C followed by washing steps. After reverse cross-linking, ChIP-DNA fragments were purified and constructed into libraries by using the ThruPLEX DNA-seq Kit (Rubicon Genomics, Takara, Japan). Amplified libraries around 300 – 500 bp were isolated from agarose gel prior to sequencing. Sequencing was performed on an Illumina Hiseq X10 platform and 150-bp paired-end reads were generated according to Illumina’s protocol. ChIP-seq data were submitted to the GEO database.

### Identification of H3K27ac regions and peaks

ChIP-seq reads were mapped to the rat genome (rn6) using Bowtie aliger allowing up to two mismatches (Langmead, 2010). Only the uniquely mapped reads were used for further analysis. SICER call peak program was used to call the peaks with a window size of 200 bp and a gap size of 400 bp, and an E value of 1. The EdgeR3 program was used to identify the differential peaks between treatment and control groups (cutoff: FC>2 and FDR<0.01).

### ChIP-qPCR assay

Four gene regions relevant to mitochondria and neurological function were selected for ChIP-qPCR assay and primers are listed in Table2. All qPCR reactions were performed on an Applied Biosystems StepOne Real-time PCR system (Thermo Fisher Scientific, USA) using iTaq Universal SYBR Green Supermix (Bio-Rad, USA) with two technical replicates. The amplification procedure was as follows: 95°C for 5 min, followed by 40 cycles of 95°C for 10 sec and 60°C for 20 sec. Relative quantification of target gene regions was performed by normalizing the Ct of samples to the percentage of Input DNA.

### Enrichment analysis of transcription factor motifs

The HOMER ChIP-seq pipeline was applied for analyzing motif enrichment (Heinz et al., 2010). Motif models were established from the TRANSFAC vertebrate database (Matys et al., 2006) and the enrichment analysis was performed on rotenone-upregulated and TFAM-KO-upregulated enhancer peaks with all peaks as the background. Enriched motifs were classified into known motifs and *de novo* motifs.

### Statistical Analysis

Data analysis was performed using GraphPad Prism 7.0. Normally distributed raw data were analyzed with either Student’s t test (2-group comparisons) or one-way ANOVA (>2-group comparisons) with Tukey post hoc test unless otherwise mentioned. Statistically significant differences were denoted as * p≤0.05, ** p<0.01, and *** p<0.001.

## Acknowledgements

This work was supported by National Institutes of Health (NIH) grants (R01 ES027245, ES026892 and NS100090 to AGK and NIH R01 NS088206 to AK). The authors acknowledge Gary Zenitsky for his assistance in editing this manuscript. Experimental disposals and salaries were also partially supported by NIH grants (R01ES25761, U01ES026721 Opportunity Fund, and R21ES028351) and Johns Hopkins Catalyst Award to ZW.

## Author contributions

A.G., Z.W., and A.G.K. jointly conceived this project and supervised the experiments. M.H., A.K., Z.W. and A.G.K. designed research. M.H., D.L., and A.C. performed experiments. M.H. and H.J. analyzed experimental results. D.K. accomplished the bioinformatic analyses. M.H., D.L., D.K., V.A., H.J., A.K., Z.W., and A.G.K. prepared manuscript. Z.W. and A.G.K. made the final call (data interpretation and presentation) on epigenomic and animal analyses, respectively.

## Reference

Aruga, J., Yokota, N., and Mikoshiba, K. (2003). Human SLITRK family genes: genomic organization and expression profiling in normal brain and brain tumor tissue. Gene 315, 87–94.

Baeza, J., Dowell, J.A., Smallegan, M.J., Fan, J., Amador-Noguez, D., Khan, Z., and Denu, J.M. (2014). Stoichiometry of site-specific lysine acetylation in an entire proteome. J Biol Chem 289, 21326–21338.

Betarbet, R., Canet-Aviles, R.M., Sherer, T.B., Mastroberardino, P.G., McLendon, C., Kim, J.H., Lund, S., Na, H.M., Taylor, G., Bence, N.F., et al. (2006). Intersecting pathways to neurodegeneration in Parkinson’s disease: effects of the pesticide rotenone on DJ-1, alpha-synuclein, and the ubiquitin-proteasome system. Neurobiol Dis 22, 404–420.

Betarbet, R., Sherer, T.B., MacKenzie, G., Garcia-Osuna, M., Panov, A.V., and Greenamyre, J.T. (2000). Chronic systemic pesticide exposure reproduces features of Parkinson’s disease. Nat Neurosci 3, 1301–1306.

Borggrefe, T., and Oswald, F. (2009). The Notch signaling pathway: transcriptional regulation at Notch target genes. Cellular and molecular life sciences: CMLS 66, 1631–1646.

Borrageiro, G., Haylett, W., Seedat, S., Kuivaniemi, H., and Bardien, S. (2018). A review of genome-wide transcriptomics studies in Parkinson’s disease. The European journal of neuroscience 47, 1–16.

Charli, A., Jin, H., Anantharam, V., Kanthasamy, A., and Kanthasamy, A.G. (2016). Alterations in mitochondrial dynamics induced by tebufenpyrad and pyridaben in a dopaminergic neuronal cell culture model. Neurotoxicology 53, 302–313.

Choudhary, C., Weinert, B.T., Nishida, Y., Verdin, E., and Mann, M. (2014). The growing landscape of lysine acetylation links metabolism and cell signalling. Nat Rev Mol Cell Biol 15, 536–550.

Creyghton, M.P., Cheng, A.W., Welstead, G.G., Kooistra, T., Carey, B.W., Steine, E.J., Hanna, J., Lodato, M.A., Frampton, G.M., Sharp, P.A., et al. (2010). Histone H3K27ac separates active from poised enhancers and predicts developmental state. Proc Natl Acad Sci U S A 107, 21931–21936.

Curry, D.W., Stutz, B., Andrews, Z.B., and Elsworth, J.D. (2018). Targeting AMPK Signaling as a Neuroprotective Strategy in Parkinson’s Disease. J Parkinsons Dis 8, 161–181.

Dawson, T.M., Ko, H.S., and Dawson, V.L. (2010). Genetic animal models of Parkinson’s disease. Neuron 66, 646–661.

Dong, Y., and Simske, J.S. (2016). Vertebrate Claudin/PMP22/EMP22/MP20 family protein TMEM47 regulates epithelial cell junction maturation and morphogenesis. Dev Dyn 245, 653–666.

Ernst, J., Kheradpour, P., Mikkelsen, T.S., Shoresh, N., Ward, L.D., Epstein, C.B., Zhang, X., Wang, L., Issner, R., Coyne, M., et al. (2011). Mapping and analysis of chromatin state dynamics in nine human cell types. Nature 473, 43–49.

Ficazzola, M.A., Fraiman, M., Gitlin, J., Woo, K., Melamed, J., Rubin, M.A., and Walden, P.D. (2001). Antiproliferative B cell translocation gene 2 protein is down-regulated post-transcriptionally as an early event in prostate carcinogenesis. Carcinogenesis 22, 1271–1279.

Fu, S., Wang, Q., Moore, J.E., Purcaro, M.J., Pratt, H.E., Fan, K., Gu, C., Jiang, C., Zhu, R., Kundaje, A., et al. (2018). Differential analysis of chromatin accessibility and histone modifications for predicting mouse developmental enhancers. Nucleic Acids Res 46, 11184–11201.

Garcia, B.A., Hake, S.B., Diaz, R.L., Kauer, M., Morris, S.A., Recht, J., Shabanowitz, J., Mishra, N., Strahl, B.D., Allis, C.D., et al. (2007). Organismal differences in post-translational modifications in histones H3 and H4. J Biol Chem 282, 7641–7655.

Goldman, S.M. (2014). Environmental toxins and Parkinson’s disease. Annu Rev Pharmacol Toxicol 54, 141–164.

Gonzalez-Reyes, L.E., Verbitsky, M., Blesa, J., Jackson-Lewis, V., Paredes, D., Tillack, K., Phani, S., Kramer, E.R., Przedborski, S., and Kottmann, A.H. (2012). Sonic hedgehog maintains cellular and neurochemical homeostasis in the adult nigrostriatal circuit. Neuron 75, 306–319.

Gordon, R., Neal, M.L., Luo, J., Langley, M.R., Harischandra, D.S., Panicker, N., Charli, A., Jin, H., Anantharam, V., Woodruff, T.M., et al. (2016). Prokineticin-2 upregulation during neuronal injury mediates a compensatory protective response against dopaminergic neuronal degeneration. Nat Commun 7, 12932.

Haley, M.J., Brough, D., Quintin, J., and Allan, S.M. (2019). Microglial Priming as Trained Immunity in the Brain. Neuroscience 405, 47–54.

Harrison, I.F., Smith, A.D., and Dexter, D.T. (2018). Pathological histone acetylation in Parkinson’s disease: Neuroprotection and inhibition of microglial activation through SIRT 2 inhibition. Neurosci Lett 666, 48–57.

Hatcher, J.M., Pennell, K.D., and Miller, G.W. (2008). Parkinson’s disease and pesticides: a toxicological perspective. Trends Pharmacol Sci 29, 322–329.

Heintzman, N.D., Hon, G.C., Hawkins, R.D., Kheradpour, P., Stark, A., Harp, L.F., Ye, Z., Lee, L.K., Stuart, R.K., Ching, C.W., et al. (2009). Histone modifications at human enhancers reflect global cell-type-specific gene expression. Nature 459, 108–112.

Heinz, S., Benner, C., Spann, N., Bertolino, E., Lin, Y.C., Laslo, P., Cheng, J.X., Murre, C., Singh, H., and Glass, C.K. (2010). Simple combinations of lineage-determining transcription factors prime cis-regulatory elements required for macrophage and B cell identities. Molecular cell 38, 576–589.

Hin, N., Newman, M., Kaslin, J., Douek, A.M., Lumsden, A., Zhou, X.-F., Mañucat-Tan, N.B., Ludington, A., Adelson, D.L., Pederson, S., et al. (2018). Accelerated brain aging towards transcriptional inversion in a zebrafish model of familial Alzheimer’s disease. BioRxiv.

Hoare, S., Copland, J.A., Wood, T.G., Jeng, Y.J., Izban, M.G., and Soloff, M.S. (1999). Identification of a GABP alpha/beta binding site involved in the induction of oxytocin receptor gene expression in human breast cells, potentiation by c-Fos/c-Jun. Endocrinology 140, 2268–2279.

Horgusluoglu, E., Nudelman, K., Nho, K., and Saykin, A.J. (2017). Adult neurogenesis and neurodegenerative diseases: A systems biology perspective. Am J Med Genet B Neuropsychiatr Genet 174, 93–112.

Huang da, W., Sherman, B.T., and Lempicki, R.A. (2009). Bioinformatics enrichment tools: paths toward the comprehensive functional analysis of large gene lists. Nucleic Acids Res 37, 1–13.

Kanehisa, M., and Goto, S. (2000). KEGG: kyoto encyclopedia of genes and genomes. Nucleic Acids Res 28, 27–30.

Kaul, S., Kanthasamy, A., Kitazawa, M., Anantharam, V., and Kanthasamy, A.G. (2003). Caspase-3 dependent proteolytic activation of protein kinase C delta mediates and regulates 1-methyl-4-phenylpyridinium (MPP+)-induced apoptotic cell death in dopaminergic cells: relevance to oxidative stress in dopaminergic degeneration. The European journal of neuroscience 18, 1387–1401.

Kim, J.M., Lee, K.H., Jeon, Y.J., Oh, J.H., Jeong, S.Y., Song, I.S., Kim, J.M., Lee, D.S., and Kim, N.S. (2006). Identification of genes related to Parkinson’s disease using expressed sequence tags. DNA research: an international journal for rapid publication of reports on genes and genomes 13, 275–286.

Kondru, N., Manne, S., Greenlee, J., West Greenlee, H., Anantharam, V., Halbur, P., and Kanthasamy, A. (2017). Integrated Organotypic Slice Cultures and RT-QuIC (OSCAR) Assay: Implications for Translational Discovery in Protein Misfolding Diseases. Sci Rep 7, 43155.

Kopp, A. (2012). Dmrt genes in the development and evolution of sexual dimorphism. Trends Genet 28, 175–184.

Kubota, K., Kent, L.N., Rumi, M.A., Roby, K.F., and Soares, M.J. (2015). Dynamic Regulation of AP-1 Transcriptional Complexes Directs Trophoblast Differentiation. Molecular and cellular biology 35, 3163–3177.

Kumar, V., Muratani, M., Rayan, N.A., Kraus, P., Lufkin, T., Ng, H.H., and Prabhakar, S. (2013). Uniform, optimal signal processing of mapped deep-sequencing data. Nature biotechnology 31, 615–622.

Langley, M.R., Ghaisas, S., Ay, M., Luo, J., Palanisamy, B.N., Jin, H., Anantharam, V., Kanthasamy, A., and Kanthasamy, A.G. (2018). Manganese exposure exacerbates progressive motor deficits and neurodegeneration in the MitoPark mouse model of Parkinson’s disease: Relevance to gene and environment interactions in metal neurotoxicity. Neurotoxicology 64, 240–255.

Li, N., Ragheb, K., Lawler, G., Sturgis, J., Rajwa, B., Melendez, J.A., and Robinson, J.P. (2003). Mitochondrial complex I inhibitor rotenone induces apoptosis through enhancing mitochondrial reactive oxygen species production. J Biol Chem 278, 8516–8525.

Li, Y.I., Wong, G., Humphrey, J., and Raj, T. (2019). Prioritizing Parkinson’s disease genes using population-scale transcriptomic data. Nature communications 10, 994.

Li, Z., Dai, H., Martos, S.N., Xu, B., Gao, Y., Li, T., Zhu, G., Schones, D.E., and Wang, Z. (2015). Distinct roles of DNMT1-dependent and DNMT1-independent methylation patterns in the genome of mouse embryonic stem cells. Genome biology 16, 115.

Manna, P.R., Dyson, M.T., Eubank, D.W., Clark, B.J., Lalli, E., Sassone-Corsi, P., Zeleznik, A.J., and Stocco, D.M. (2002). Regulation of steroidogenesis and the steroidogenic acute regulatory protein by a member of the cAMP response-element binding protein family. Molecular endocrinology 16, 184–199.

Martos, S.N., Li, T., Ramos, R.B., Lou, D., Dai, H., Xu, J.C., Gao, G., Gao, Y., Wang, Q., An, C., et al. (2017). Two approaches reveal a new paradigm of ‘switchable or genetics-influenced allele-specific DNA methylation’ with potential in human disease. Cell Discov 3, 17038.

Marzi, S.J., Leung, S.K., Ribarska, T., Hannon, E., Smith, A.R., Pishva, E., Poschmann, J., Moore, K., Troakes, C., Al-Sarraj, S., et al. (2018). A histone acetylome-wide association study of Alzheimer’s disease identifies disease-associated H3K27ac differences in the entorhinal cortex. Nat Neurosci 21, 1618–1627.

Matilainen, O., Quirós, P.M., and Auwerx, J. (2017). Mitochondria and Epigenetics - Crosstalk in Homeostasis and Stress. Trends Cell Biol 27, 453–463.

Matys, V., Kel-Margoulis, O.V., Fricke, E., Liebich, I., Land, S., Barre-Dirrie, A., Reuter, I., Chekmenev, D., Krull, M., Hornischer, K., et al. (2006). TRANSFAC and its module TRANSCompel: transcriptional gene regulation in eukaryotes. Nucleic Acids Res 34, D108–110.

Maurano, M.T., Humbert, R., Rynes, E., Thurman, R.E., Haugen, E., Wang, H., Reynolds, A.P., Sandstrom, R., Qu, H., Brody, J., et al. (2012). Systematic localization of common disease-associated variation in regulatory DNA. Science (New York, NY) 337, 1190–1195.

Miyazaki, T., Iwasawa, M., Nakashima, T., Mori, S., Shigemoto, K., Nakamura, H., Katagiri, H., Takayanagi, H., and Tanaka, S. (2012). Intracellular and extracellular ATP coordinately regulate the inverse correlation between osteoclast survival and bone resorption. J Biol Chem 287, 37808–37823.

Ostuni, R., Piccolo, V., Barozzi, I., Polletti, S., Termanini, A., Bonifacio, S., Curina, A., Prosperini, E., Ghisletti, S., and Natoli, G. (2013). Latent enhancers activated by stimulation in differentiated cells. Cell 152, 157–171.

Panicker, N., Sarkar, S., Harischandra, D.S., Neal, M., Kam, T.I., Jin, H., Saminathan, H., Langley, M., Charli, A., Samidurai, M., et al. (2019). Fyn kinase regulates misfolded α-synuclein uptake and NLRP3 inflammasome activation in microglia. J Exp Med 216, 1411–1430.

Plagge, A., Kelsey, G., and Germain-Lee, E.L. (2008). Physiological functions of the imprinted Gnas locus and its protein variants Galpha(s) and XLalpha(s) in human and mouse. J Endocrinol 196, 193–214.

Polymeropoulos, M.H., Lavedan, C., Leroy, E., Ide, S.E., Dehejia, A., Dutra, A., Pike, B., Root, H., Rubenstein, J., Boyer, R., et al. (1997). Mutation in the alpha-synuclein gene identified in families with Parkinson’s disease. Science 276, 2045–2047.

Quirós, P.M., Mottis, A., and Auwerx, J. (2016). Mitonuclear communication in homeostasis and stress. Nat Rev Mol Cell Biol 17, 213–226.

Seo, J., Ottesen, E.W., and Singh, R.N. (2014). Antisense methods to modulate pre-mRNA splicing. Methods Mol Biol 1126, 271–283.

Siani, F., Greco, R., Levandis, G., Ghezzi, C., Daviddi, F., Demartini, C., Vegeto, E., Fuzzati-Armentero, M.T., and Blandini, F. (2017). Influence of Estrogen Modulation on Glia Activation in a Murine Model of Parkinson’s Disease. Frontiers in neuroscience 11, 306.

Sodersten, E., Feyder, M., Lerdrup, M., Gomes, A.L., Kryh, H., Spigolon, G., Caboche, J., Fisone, G., and Hansen, K. (2014). Dopamine signaling leads to loss of Polycomb repression and aberrant gene activation in experimental parkinsonism. PLoS genetics 10, e1004574.

Song, C., Kanthasamy, A., Anantharam, V., Sun, F., and Kanthasamy, A.G. (2010). Environmental neurotoxic pesticide increases histone acetylation to promote apoptosis in dopaminergic neuronal cells: relevance to epigenetic mechanisms of neurodegeneration. Molecular pharmacology 77, 621–632.

Stephano, F., Nolte, S., Hoffmann, J., El-Kholy, S., von Frieling, J., Bruchhaus, I., Fink, C., and Roeder, T. (2018). Impaired Wnt signaling in dopamine containing neurons is associated with pathogenesis in a rotenone triggered Drosophila Parkinson’s disease model. Scientific reports 8, 2372.

Sun, W., Poschmann, J., Cruz-Herrera Del Rosario, R., Parikshak, N.N., Hajan, H.S., Kumar, V., Ramasamy, R., Belgard, T.G., Elanggovan, B., Wong, C.C.Y., et al. (2016). Histone Acetylome-wide Association Study of Autism Spectrum Disorder. Cell 167, 1385–1397 e1311.

Svinkina, T., Gu, H., Silva, J.C., Mertins, P., Qiao, J., Fereshetian, S., Jaffe, J.D., Kuhn, E., Udeshi, N.D., and Carr, S.A. (2015). Deep, Quantitative Coverage of the Lysine Acetylome Using Novel Anti-acetyl-lysine Antibodies and an Optimized Proteomic Workflow. Molecular & cellular proteomics: MCP 14, 2429–2440.

Sánchez-Flores, M., Pásaro, E., Bonassi, S., Laffon, B., and Valdiglesias, V. (2015). γH2AX assay as DNA damage biomarker for human population studies: defining experimental conditions. Toxicol Sci 144, 406–413.

Tanner, C.M., Kamel, F., Ross, G.W., Hoppin, J.A., Goldman, S.M., Korell, M., Marras, C., Bhudhikanok, G.S., Kasten, M., Chade, A.R., et al. (2011). Rotenone, paraquat, and Parkinson’s disease. Environmental health perspectives 119, 866–872.

Tiwari, P.C., and Pal, R. (2017). The potential role of neuroinflammation and transcription factors in Parkinson disease. Dialogues in clinical neuroscience 19, 71–80.

Trancikova, A., Tsika, E., and Moore, D.J. (2012). Mitochondrial dysfunction in genetic animal models of Parkinson’s disease. Antioxid Redox Signal 16, 896–919.

Utami, K.H., Hillmer, A.M., Aksoy, I., Chew, E.G., Teo, A.S., Zhang, Z., Lee, C.W., Chen, P.J., Seng, C.C., Ariyaratne, P.N., et al. (2014). Detection of chromosomal breakpoints in patients with developmental delay and speech disorders. PloS one 9, e90852.

Valentine, M.N.Z., Hashimoto, K., Fukuhara, T., Saiki, S., Ishikawa, K.I., Hattori, N., and Carninci, P. (2019). Multi-year whole-blood transcriptome data for the study of onset and progression of Parkinson’s Disease. Scientific data 6, 20.

van der Werf, I.M., Van Dijck, A., Reyniers, E., Helsmoortel, C., Kumar, A.A., Kalscheuer, V.M., de Brouwer, A.P., Kleefstra, T., van Bokhoven, H., Mortier, G., et al. (2017). Mutations in two large pedigrees highlight the role of ZNF711 in X-linked intellectual disability. Gene 605, 92–98.

Vlahopoulos, S.A., Logotheti, S., Mikas, D., Giarika, A., Gorgoulis, V., and Zoumpourlis, V. (2008). The role of ATF-2 in oncogenesis. BioEssays: news and reviews in molecular, cellular and developmental biology 30, 314–327.

Wang, X.F., Li, S., Chou, A.P., and Bronstein, J.M. (2006). Inhibitory effects of pesticides on proteasome activity: implication in Parkinson’s disease. Neurobiol Dis 23, 198–205.

Wang, Z., Zang, C., Cui, K., Schones, D.E., Barski, A., Peng, W., and Zhao, K. (2009). Genome-wide mapping of HATs and HDACs reveals distinct functions in active and inactive genes. Cell 138, 1019–1031.

Wang, Z., Zang, C., Rosenfeld, J.A., Schones, D.E., Barski, A., Cuddapah, S., Cui, K., Roh, T.Y., Peng, W., Zhang, M.Q., et al. (2008). Combinatorial patterns of histone acetylations and methylations in the human genome. Nat Genet 40, 897–903.

Weinert, B.T., Iesmantavicius, V., Moustafa, T., Schölz, C., Wagner, S.A., Magnes, C., Zechner, R., and Choudhary, C. (2014). Acetylation dynamics and stoichiometry in Saccharomyces cerevisiae. Mol Syst Biol 10, 716.

Wu, S., Lei, L., Song, Y., Liu, M., Lu, S., Lou, D., Shi, Y., Wang, Z., and He, D. (2018). Mutation of hop-1 and pink-1 attenuates vulnerability of neurotoxicity in C. elegans: the role of mitochondria-associated membrane proteins in Parkinsonism. Exp Neurol 309, 67–78.

Xia, P., Wang, S., Huang, G., Du, Y., Zhu, P., Li, M., and Fan, Z. (2014). RNF2 is recruited by WASH to ubiquitinate AMBRA1 leading to downregulation of autophagy. Cell Res 24, 943–958.

Zanon, A., Pramstaller, P.P., Hicks, A.A., and Pichler, I. (2018). Environmental and Genetic Variables Influencing Mitochondrial Health and Parkinson’s Disease Penetrance. Parkinsons Dis 2018, 8684906.

Zhang, X., Hamada, J., Nishimoto, A., Takahashi, Y., Murai, T., Tada, M., and Moriuchi, T. (2007). HOXC6 and HOXC11 increase transcription of S100beta gene in BrdU-induced in vitro differentiation of GOTO neuroblastoma cells into Schwannian cells. Journal of cellular and molecular medicine 11, 299–306.

