## Supplemental Figure S1~S9 for "Mitochondrial Dysfunction Induces Epigenetic Dysregulation by H3K27 Hyperacetylation to Perturb Active Enhancers in Parkinson’s Disease Models"

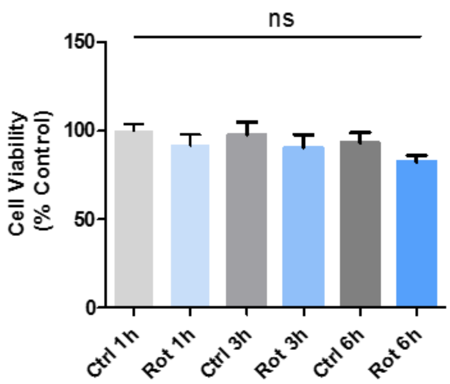


**Figure S1**. MTS assay of cell viability after treating N27 cells with the mitochondrial complex-1 inhibitor rotenone (1 μM) for 1, 3, and 6 h. Data analyzed via one-way ANOVA with Tukey's post hoc test (ns, not significant) and are represented as mean ± s.e.m with n=8.

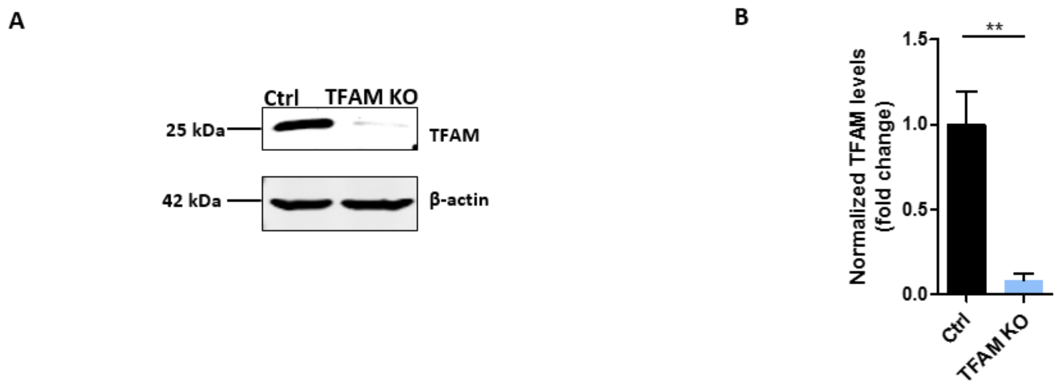


**Figure S2**. Mitochondrial transcription factor A (TFAM) was knocked out in N27 DAergic neuronal cells. (A) Representative immunoblot for TFAM. (B) Densitometric analysis of bands in immunoblot. Bar graph shows mean ± s.e.m of unpaired two-tailed t tests with n=4. $*p \leq 0.05$, $**p < 0.01$.

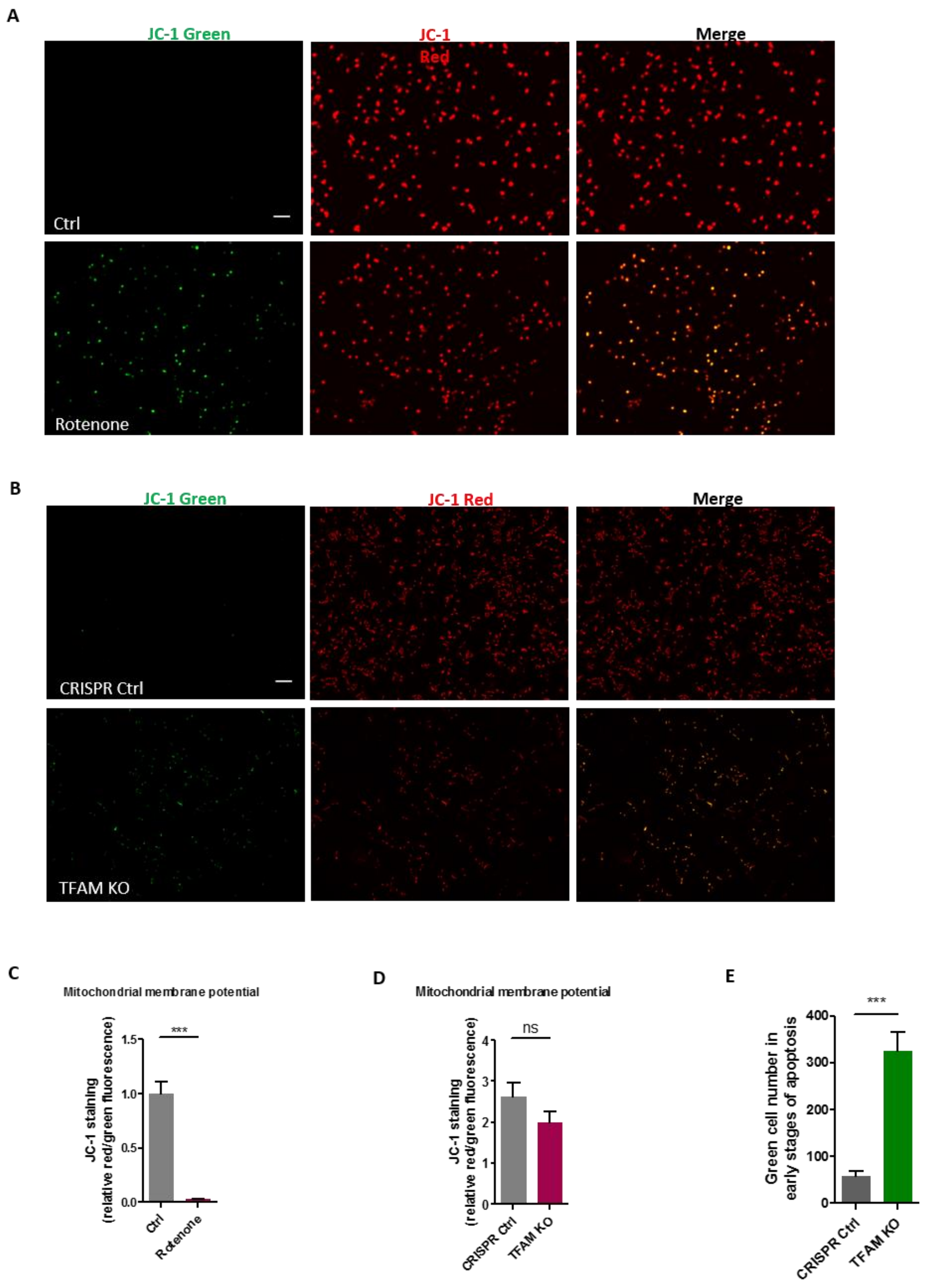


**Figure S3**: Increased depolarization of mitochondrial membrane potential in (A) rotentone-exposed and (B) TFAM-KO N27 cells as measured by JC1 probe. Scale bars, 100 μm. Quantitative analysis of the $\Delta\psi m$ by the ratio of the red and green fluorescence for (C) rotenone-exposed and (D) TFAM-KO N27 cells. (E) Cell number of TFAM KO that shows in green color. Experiments were repeated three times with 3-4 samples per time. Bar graphs show mean ± s.e.m of unpaired two-tailed t tests. ns, not significant; $*p \leq 0.05$; $**p<0.01$; $***p<0.001$.

A

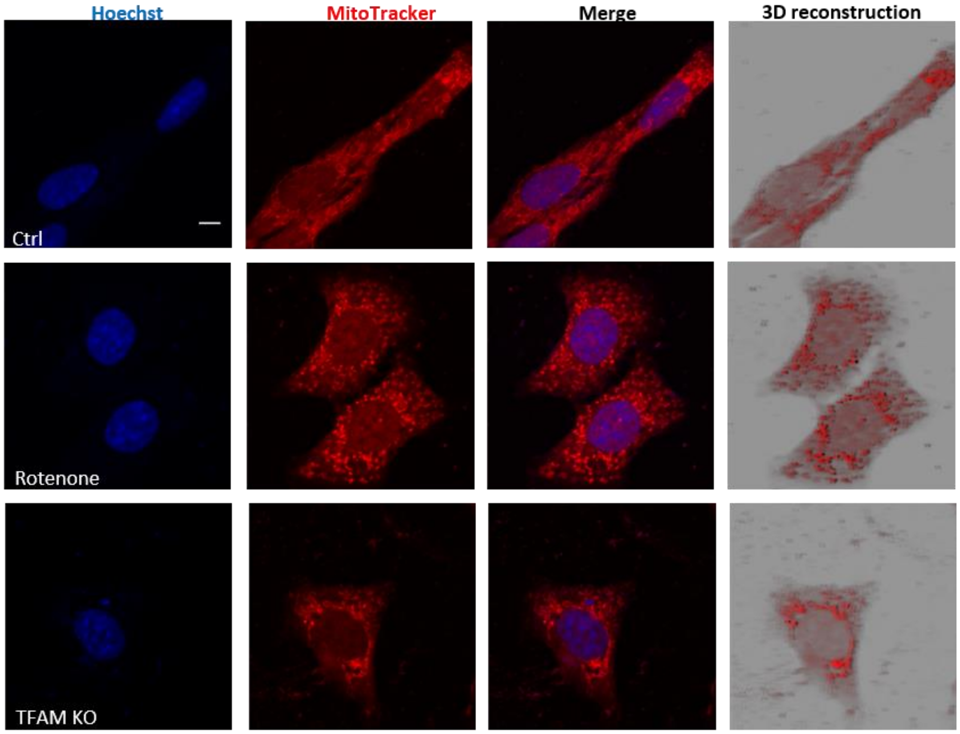


B

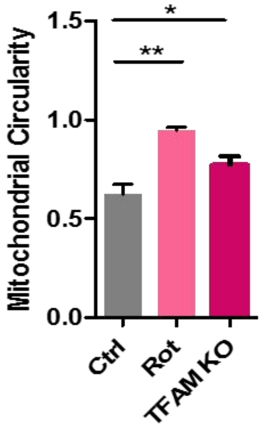


**Figure S4**: Mitochondrial morphological changes in rotentone-exposed and TFAM KO N27 neuronal cells. (A) Mitochondrial structure measurement by MitoTracker (red) dye showing induced small, punctuate mitochondria, compared to elongated mitochondria in control. Independent experiments were repeated at least three times. Scale bars, 10 µm. (B) The quantification of mitochondrial circularity. Bar graph shows mean ± s.e.m of unpaired two-tailed t tests. ns, not significant; *$p \leq 0.05$; **$p < 0.01$; ***$p < 0.001$.

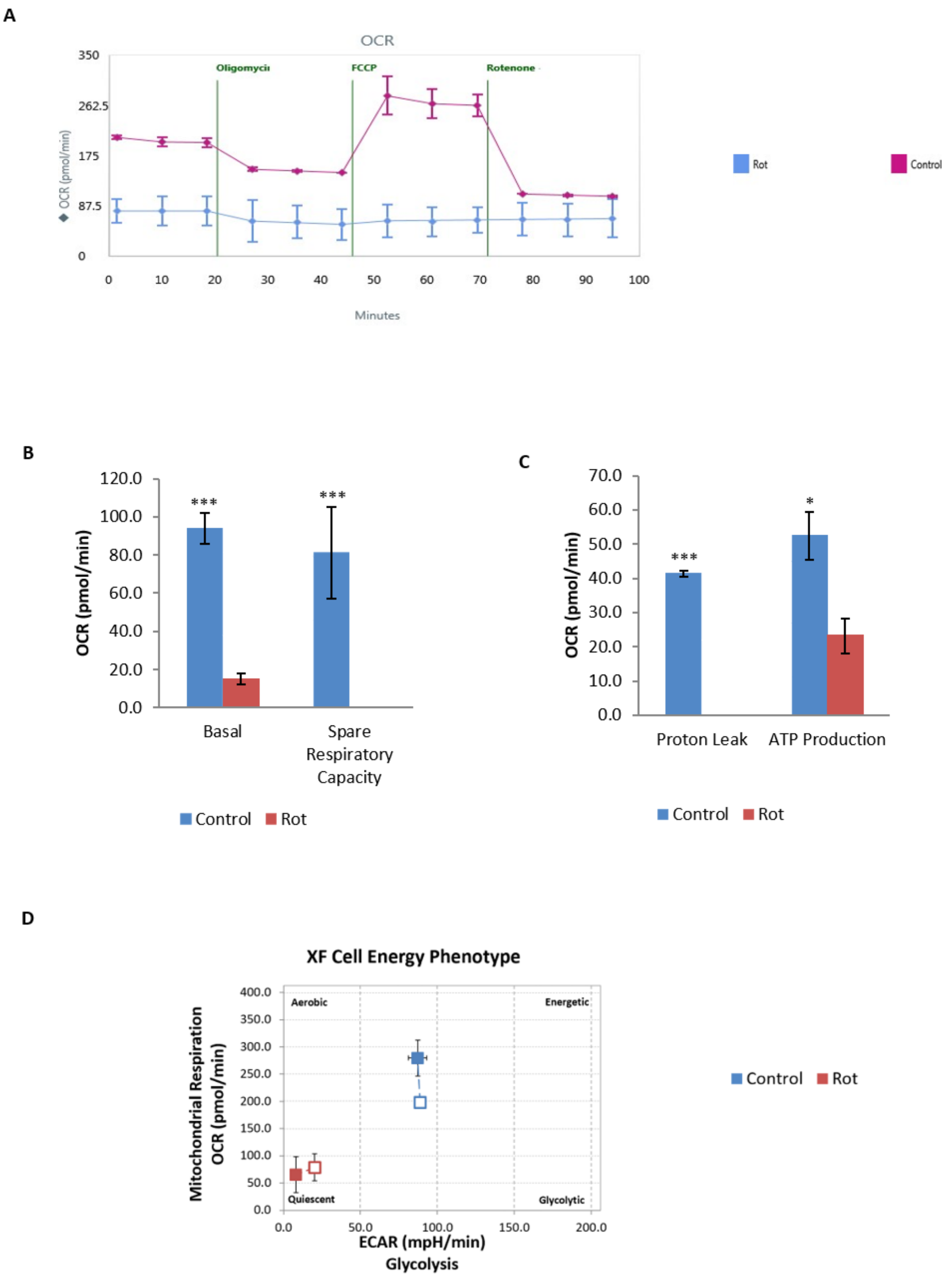


**Figure S5**. Impaired mitochondrial bioenergetics of rotenone-exposed N27 neuronal cells revealed by XFe24 Seahorse mitochondrial stress test. (A, B, and C) Respiration plot, basal respiration, spare respiratory capacity, proton leak, and ATP production. (D) Cellular phenotype plot comparing OCR on the y-axis and ECAR on the x-axis. Three biological replicates for each treatment. Bar graphs show mean ± s.e.m of unpaired two-tailed t tests. *p≤0.05, and **p<0.01.

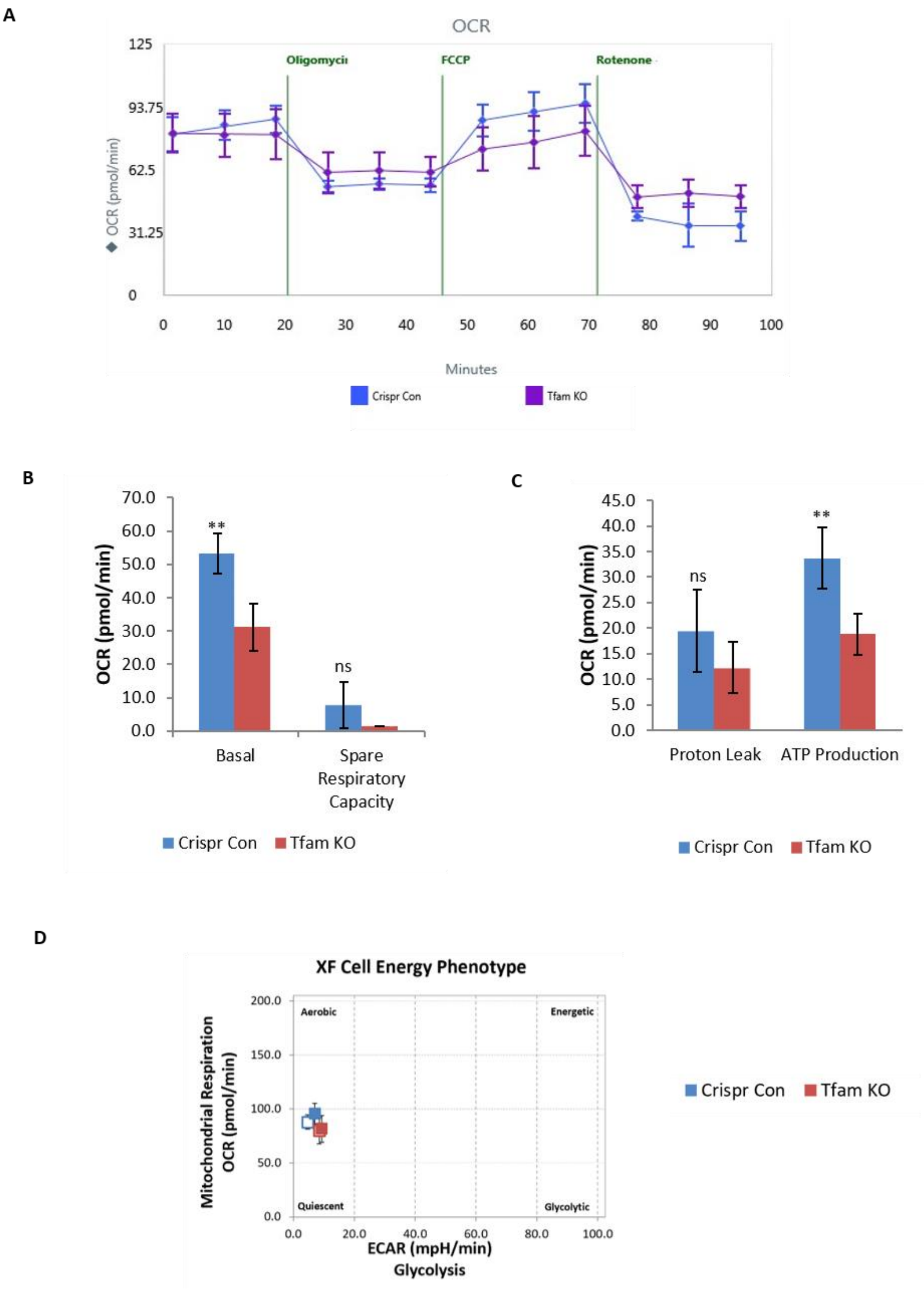


**Figure S6**. Impaired mitochondrial bioenergetics of TFAM-knockout N27 neuronal cells revealed by XFe24 Seahorse mitochondrial stress test. (A, B and C) Respiration plot, basal respiration, spare respiratory capacity, proton leak, and ATP production. (D) Cellular phenotype plot comparing OCR on the y-axis and ECAR on the x-axis. Five independent measurements for each data point. Bar graphs show mean ± s.e.m of unpaired two-tailed t tests. ns, not significant; $*p \leq 0.05$, and $**p < 0.01$.

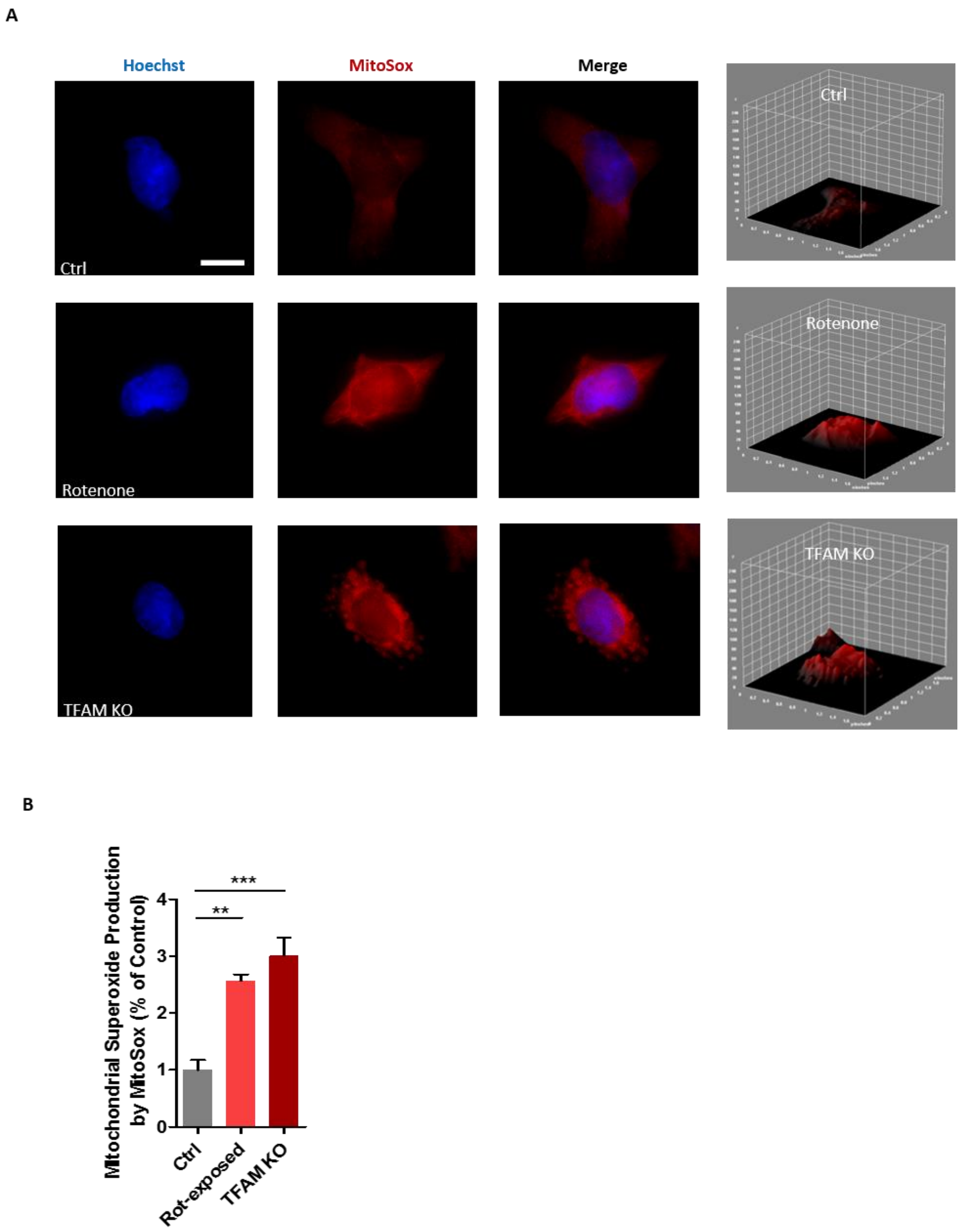


**Figure S7**: Oxidative stress induced in rotentone-exposed and TFAM-KO N27 neuronal cells. (A) MitoSox (red) assay showing mitoROS production coupled with 3D plots of MitoSox fluorescent intensity. Independent experiments were repeated at least four times. Scale bars, 10 µm. (B) Quantification of mitochondrial superoxide production. Error bars represent mean±s.e.m. of one-way ANOVA followed by Tukey's post hoc test. *p≤0.05; **p<0.01; ***p<0.001.

A

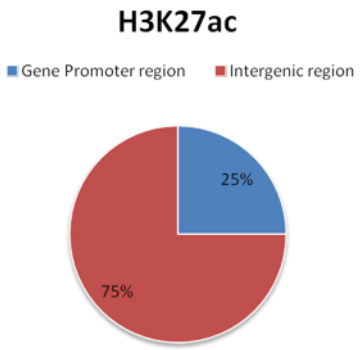


B

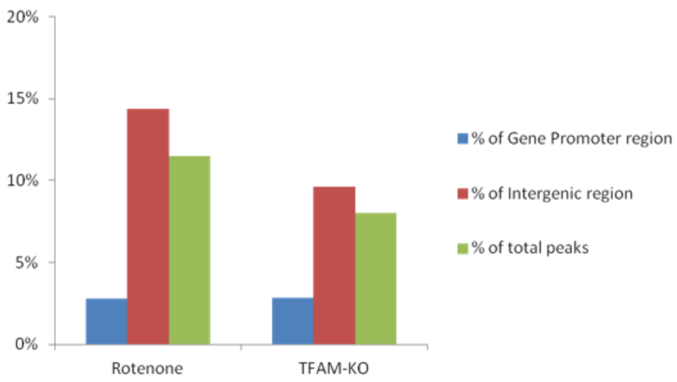


**Figure S8**. (A) Genome-wide H3K27ac distribution in N27 cells. Approximately 25% of the identified H3K27ac peaks are located at annotated genes or their proximal promoters and 75% of the peaks correspond to the intergenic regions (TSS±2.5kb). (B) Genome-wide H3K27ac modification alterations induced by mitochondrial impairment. Approximately 11.5% of H3K27ac peaks were significantly changed after rotenone treatment, with 2.79% and 14.36% located at annotated genes (proximal promoters) and distal promoters, respectively. Upon TFAM knockout, 8.0% of H3K27ac peaks were significantly changed, with 2.82% and 9.63% at annotated genes (proximal promoters) and distal promoters, respectively.

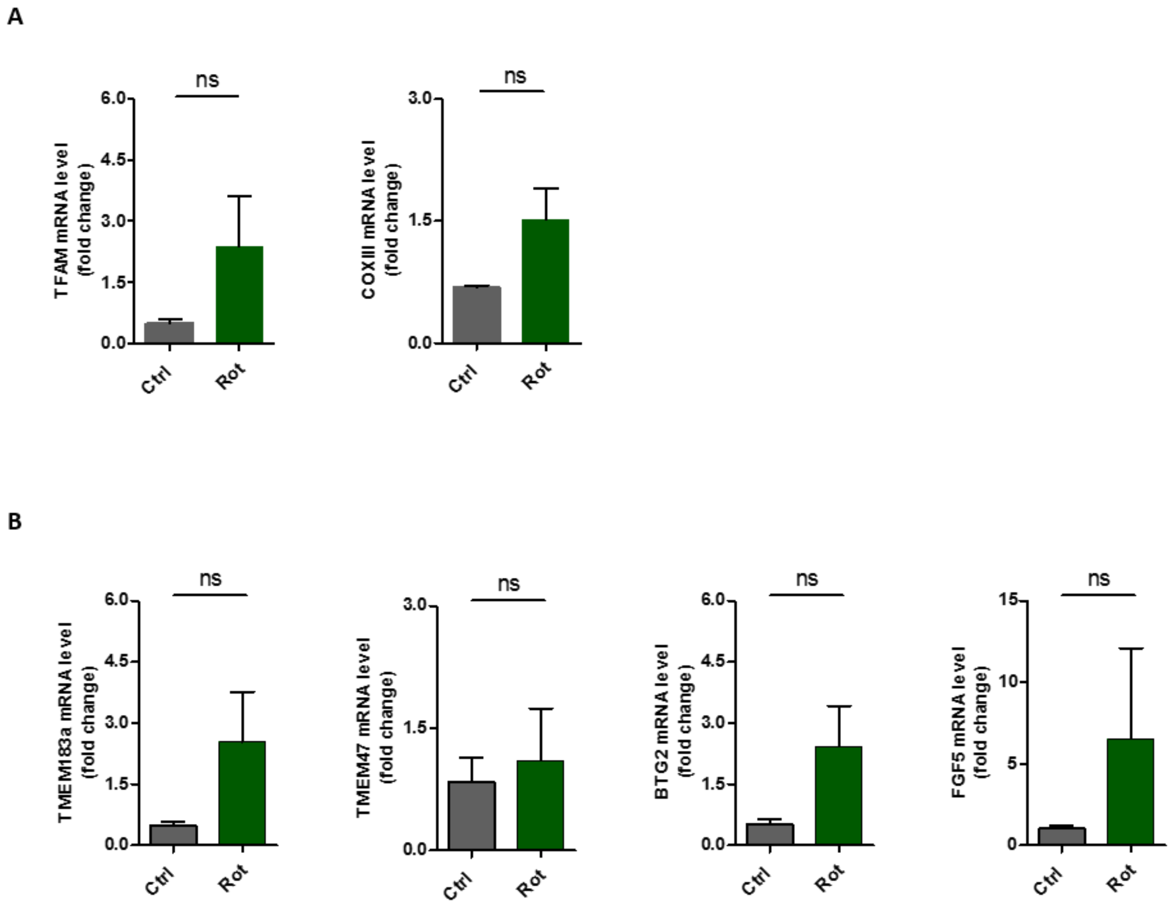


**Figure S9**. Increased mRNA levels of selected genes in rotenone-exposed midbrain slices from wild-type C57BL/6 mice. (A) qRT-PCR validation of mitochondrial impairment using TFAM and COXIII as biomarkers in midbrain slices acutely treated with either 25 µM rotenone or vehicle (DMSO) for 3 h. (B) qRT-PCR analysis of mouse midbrain slices dissected from 7- to 12-day-old pups. Four differentially expressed genes related to mitochondrial or neuronal functions were selected from the upregulated overlapping gene pool attained by integrating RNA-seq and ChIP-seq analysis. At least four independent experiments were measured. Bar graphs show mean ± s.e.m of unpaired two-tailed t tests. ns, not significant; *p≤0.05.
